# Insight Into the Molecular Parameters of PEI Promoting an Efficient Gene Delivery into Cells

**DOI:** 10.1101/2024.11.01.621598

**Authors:** Paulina Alejandra Montaño González, Sylvain Tranchimand, Manon Rochedy, Patricia Leroyer, Jean-Paul Chapel, Soizic Chevance, Christophe Schatz, Fabienne Gauffre, Pascal Loyer, Lourdes Mónica Bravo Anaya

## Abstract

The development of natural or synthetic polycations able to interact with nucleic acids and condense them into nanoparticles known as polyplexes, faces several unresolved challenges at the cellular level. Key issues include the intracellular trafficking of polyplexes, the endosomal escape and the release of nucleic acids into the cytosol, which are considered major bottlenecks for efficient protein expression. Here, we aim at gaining fundamental insights into the stability of polyplexes in biological media and their uptake and intracellular trafficking, while correlating data of the expression of reporter protein with both the molecular characteristics of various poly(ethylenimines) (PEI) and the physicochemical characteristics of PEI/peGFP-C3 polyplexes. For this, we chosen four samples of PEI, selected as a model polycation, with different molecular weights (Mw = 0.8, 20, 25 and 60 kg/mol) and structures (linear and branched). We found that the *in vitro* and *in vivo* stability of PEI/peGFP-C3 polyplexes, their cell internalization and transfection efficiency is dependent on the variation of polycation Mw and structure, as well as the intrinsic properties of polyplexes, such as the charge ratio (R=[N^+^]/[P^−^]). A relation between the percentage of positive cells to green fluorescent protein (GFP) and the amount of internalized nucleic acid (cyanine 5-peGFP-C3) allowed revealing the molecular characteristics of PEI promoting both higher both cell internalization and GFP expression on HEK293T cells. In the long term, the outcome of this work will be to propose guidelines to help design more effective, and less cytotoxic non-viral gene carriers with a great potential for new therapeutic applications.

## 1. Introduction

Nucleic acid-based therapies are groundbreaking and effective treatments that employ therapeutic nucleic acids like DNA, RNA, and oligonucleotides to transiently or stably modify gene expression [1–3]. This approach targets various severe human diseases, including cancers, cystic fibrosis, immunodeficiency, and cardiovascular conditions [4–7]. Nucleic acid carriers are primarily classified into two groups: viral and nonviral carriers [8,9]. Even if viral vectors exhibit a very high transduction efficiency, concerns about medical safety and the cost-effectiveness of viral carriers have increasingly limited their use in clinical settings [10]. To address these issues, many researchers have been investigating non-viral carriers, particularly liposomes, peptides, and polycations, which offer improved safety and the possibility for large-scale production [11–13]. Polycations employed as gene delivery vectors must effectively protect genetic material from degradation such as nucleolytic enzymes, to ensure prolonged circulation in the bloodstream and targeted release in specific tissues or cells [14]. Common polycation-based vectors include poly(ethylenimine) (PEI), poly-L-lysine (PLL), poly-(2-(dimethylamino)ethyl methacrylate) (PDMAEMA), and chitosan, all of which have been extensively studied [15–19].

PEI is a well-studied gene delivery nanocarrier known for its ability to efficiently condense DNA and form stable PEI/DNA polyelectrolyte complexes, also known as polyplexes [20–23]. PEI is classified as a linear or branched polycation and is available in a broad range of molecular weights (Mw) and different degrees of branching [24]. Although PEI has demonstrated limited biodegradability, PEI/nucleic acid polyplexes have been used in several clinical trials, depicting favorable safety profiles [25,26]. The high transfection efficiency of PEI is attributed to the abundance of amine groups in its chain, that can become protonated when the endosomes acidify [27,28]. This buffering ability is believed to facilitate the release of polyplexes from the endosomes via the “proton sponge” mechanism [29–31]. However, even if numerous studies focused on understanding the mechanisms behind the uptake and intracellular transport of non-viral vectors, the mechanisms associated with PEI-mediated gene delivery are not fully understood [31]. As an example, the exact way in which PEI facilitates the endosomal escape of polyplexes is still a subject of debate [32–34]. Furthermore, it is well known that polyplexes prepared using different PEI samples and preparation protocols exhibit variable physicochemical behavior and lead to different transfection efficiencies [35–37]. For instance, the most widely used PEIs for transfection include linear PEI (*l*-PEI) with a Mw around 22 kg/mol and branched PEI (*b*-PEI) with a Mw around 25 kg/mol [38,39]. It has also been reported that the majority of internalized PEI/DNA polyplexes reach the lysosomes, while only a small percentage manage to escape from the endosomes or lysosomes [40]. This was explained with the basis that several conditions must be met to induce the proton sponge mechanism, facilitating membrane permeabilization. These conditions include the concentration of PEI, the composition and pH of the endosome [41].

In this work we aimed at studying the effect of PEI structure and Mw, as well as PEI/plasmid peGFP-C3 polyplexes charge ratio, R=[N^+^]/[P^−^]=3, 5, 10 and 15 (where [N^+^] corresponds to the molar concentration of PEI positively charged amino groups and [P^−^] corresponds to the molar concentration of negatively charged phosphate groups from DNA), on their stability in biological media (phosphate buffer saline, PBS, *i.e.* isotonic saline solution for cells, serum-free culture medium, and serum-containing culture medium), cell internalization, cell viability and transfection efficiency in HEK293T cells. The HEK293T cells were chosen because of their high transfectability, which provides a suitable cell system to assess different parameters of transfection such as the nucleic acid internalization, the expression level of a reporter protein and cell viability. Four PEI were selected to carry on this study: linear PEI (Mw = 20 kg/mol) and branched PEI (Mw= 0.8, 25 and 60 kg/mol). The plasmid peGFP-C3 was chosen to allow a straightforward detection of the expression of the green fluorescent protein (GFP) characterizing the transfection efficiency. In addition, we used the same plasmid labeled with cyanine 5 (Cy5) to monitor intracellular accumulation of plasmid and to correlate with the GFP expression levels obtained with the different polyplexes. A variety of physicochemical analyses such as dynamic light scattering (DLS), ζ-potential and small angle X-ray scattering (SAXS) combined with gel electrophoresis, fluorescence confocal microscopy and biological tools were used to characterize the polyplexes and evaluate their intracellular fate. From the obtained results, we were able to propose and highlight some correlations between physicochemical characteristics, the amount of internalized plasmid and the transfection efficiencies in HEK293T cells.

## 2. Materials and methods

### 2.1. Materials

A plasmid encoding for Green Fluorescent Protein (GFP), *i.e.* peGFP-C3 plasmid (with 4727 bp, equivalent to an average molecular weight of 3,100,912 g/mol) was purified from an *E. coli* culture using a NucleoBond PC 1000 kit following the manufacturer’s indications. peGFP-C3 purified plasmid was dissolved in deionized sterile water. An UV spectrophotometer, Nanodrop Mettler Toledo, was used to determine the purity and concentration of the plasmid. Four PEI samples with different structures and molecular weights were selected: branched PEI with 0.8 and 25 kg/mol (*b*-PEI_0.8_ and *b*-PEI_25_, respectively) were purchased from Merck (Codes 408719 and 408727, respectively), 50% w/aq. branched PEI with 60 kg/mol (*b*-PEI_60_) was purchased from Acros Organics (Code 1785771000), and linear PEI hydrochloride with 20 kg/mol (*l*-PEI_20_) was purchased from Merck (Code 764965). MIR3725 Label IT Nucleic Acid Labeling Kit, cyanine 5 (Cy5), was purchased from EUROMEDEX. NaOH in pellets with impurities ≤0.001%, anhydrous NaCl (99%) and HCl (37%) were supplied by Merck. Dulbecco’s Modified Eagle’s Medium, DMEM 1X, (11960044, Gibco) was selected for culturing human embryonic kidney cells, HEK-293T cells [42]. DMEM was supplemented with 10% fetal bovine serum (FBS), consisting of a mixture of 25% sera FetalClone II (SH30066.03, HyCloneTM) and 75% Biosera (FB-1001/500), 2 mM L-glutamine (25030-024, Gibco) and respectively 100 U/mL and 100 μg/mL of penicillin and streptomycin (15070-063, Gibco).

### 2.2. PEI solubilization and peGFP-C3 solutions preparation

*b*-PEI_0.8_, *b*-PEI_25_ and *b*-PEI_60_ solutions were prepared at a concentration of 1 mg/mL in deionized (DI) water and then adjusted with HCl 0.1 M to a pH of 7.4. *l*-PEI_20_ solutions were prepared at a concentration of 1 mg/mL in DI water and then adjusted with NaOH 0.1 M to a pH of 7.4. The purified peGFP-C3 plasmid was diluted to a concentration of 0.034 mg/mL in water in order to prepare PEI/peGFP-C3 polyplexes. Cy5-labeled peGFP-C3 plasmid was prepared following the instructions of the commercial kit supplied by EUROMEDEX. All PEI solutions were sealed with Parafilm® to prevent solvent evaporation and stored in refrigeration at 4 °C. peGFP-C3 solutions were also sealed and stored in a freezer at −20°C.

### 2.3. ζ-potential measurements

ζ-potential measurements were carried on in a Malvern Zetasizer NanoZS instrument at a temperature of 25 °C. PEI/peGFP-C3 polyplexes formation was monitored in terms of ζ-potential as a function of polyplexes charge ratio (R=[N^+^]/[P^−^]) upon the progressive addition of each PEI solution (previously adjusted at a pH of 7.4) into the peGFP-C3 solution at a concentration of 0.005 mg/mL under continuous agitation and ambient temperature. The stabilization of PEI/peGFP-C3 polyplexes was reached after 1 min after each PEI addition. A volume of 1 mL of the suspension was transferred into the Zetasizer NanoZS cell (DTS1070) for each measurement. After each ζ-potential measurement, the analyzed sample was transferred back into the initial solution to maintain a nearly constant volume of PEI/peGFP-C3 polyplexes suspension, before the next addition of PEI. The ζ-potential value was calculated through the classical Smoluchowski expression using the measured electrophoretic mobility by the Malvern instrument [43]. The Malvern instrument was set to record three runs for each measurement. Overall, two independent PEI/peGFP-C3 titrations were done. ζ-potential results are presented as the average of the three measurements with the +/− standard deviation of two independent PEI/peGFP-C3 titrations or polyplexes replicates.

### 2.4. Agarose gel electrophoresis

PEI/peGFP-C3 polyplexes stoichiometry, as well as their stability in terms of peGFP-C3 release in PBS, culture media and serum, commonly used as biological media during cell transfections, were studied by gel retardation assays. For stoichiometry experiments, PEI/peGFP-C3 polyplexes were prepared with 20 μL of a peGFP-C3 solution at a concentration of 0.005 mg/mL with increasing volumes of each PEI sample at a concentration of 0.032 mg/mL for *b-*PEI_0.8_, 0.0247 mg/mL for *l-*PEI_20_, 0.0235 mg/mL for *b-*PEI_25_ and 0.0192 mg/mL for *b-*PEI_60_, at a pH of 7.4. Then, a volume of 4 μL of 4X-loading buffer (0.1 mM EDTA, 0.05 % bromo phenol blue, 50 % glycerol) was added to the mixture. A volume of 10 μL of each PEI/peGFP-C3 polyplex sample was placed in the wells of a 0.8 % agarose gel in 1x TAE buffer (40 mM Tris, 20 mM acetic acid, 1 mM EDTA). SYBR® Safe (Life Technologies, CA, USA) was used to reveal the peGFP-C3 plasmid band. The gel electrophoresis was run at 100 V during 45 min. The Molecular Imager® Gel Doc^TM^ XR+ with Image Lab^TM^ Software was used to analyze the resulting gels.

PEI/peGFP-C3 polyplexes stability, in terms of plasmid release in presence of PBS, culture media and serum, was studied by gel electrophoresis. For this, polyplexes using the four different PEI samples were prepared with 1 mL of a peGFP-C3 solution at a concentration of 0.034 mg/mL, at R= 3 and 10. All formulated polyplexes were diluted in water to have a final concentration of peGFP-C3 of 0.022 mg/mL. In order to mimic transfection conditions, we selected the following composition: 1 % v/v PBS 1X, 81 % v/v of DMEM culture medium, 9 % v/v FBS and 9 % v/v of polyplexes. For the samples studied without either PBS, culture medium or FBS, the corresponding volumes were replaced with DI water. A volume of 4 μL of loading buffer (0.1 mM EDTA, 0.05 % bromophenol blue, 50 % glycerol) was added to 10 μL of each sample. Finally, a volume of 10 μL of each sample was deposited in the wells of a 0.8 % agarose gel in 1x TAE buffer. 10 µL of peGFP-C3 at a concentration of 0.002 mg/mL was used as a control. The gel electrophoresis was run at 100 V during 45 min and the Molecular Imager® Gel Doc^TM^ XR+ with Image Lab^TM^ Software was used to analyze the resulting gels.

### 2.5. Polyplexes preparation following the “one-shot and vortex” mixing procedure

After determination of PEI/peGFP-C3 polyplexes stoichiometry from gel electrophoresis and ζ-potential measurements, polyplexes were prepared with the four PEI samples at four selected charge ratios (R = [N^+^]/[P^−^] = 3, 5, 10 and 15). Specific volumes of each PEI solution prepared at a concentration of 1 mg/mL and adjusted at a pH of 7.4, were rapidly added to a peGFP-C3 solution with a concentration of 0.034 mg/mL, followed by fast vortexed-mixing of the suspension for 10 seconds.

### 2.6. Dynamic light scattering (DLS) measurements

PEI/peGFP-C3 polyplexes, prepared using 1 mL of a peGFP-C3 solution at a concentration of 0.034 mg/mL, were studied by DLS measurements, in a Malvern ZetaSizer Nano ZS instrument (Malvern, U.K.), equipped with a standard HeNe laser emitting at 632.8 nm with a detection angle of 173 °. After a temperature equilibration time of 2 minutes, to reach 25 °C, the correlation functions were averaged from three measurements of 2 set runs, lasting 30 sec each one. Intensity-averaged size distributions were obtained by applying the NNLS (non-negative least squares) algorithm included in the « General purpose » analysis model. The Cumulants method was also used to analyze the data of monodisperse samples and the Z-average of the hydrodynamic diameter, D_H_, was determined using the Stokes−Einstein equation, assuming spherical particles [43,44]. The values of D_H_ and PDI (polydispesity index) are presented with the ± standard deviation, SD. Two independent formulation replicates were studied.

After formulating PEI/peGFP-C3 polyplexes, their stability was studied in terms of size evolution or aggregation by DLS in the different media used during cell transfections. For this, 100 μL of the initial formulation were mixed with 900 μL of 1x PBS. The D_H_ and PDI were measured right after homogenization of the suspension and after 30 min rest. Next, 500 μL of this solution was mixed with 500 μL of DMEM culture medium; the D_H_ and PDI were measured right after homogenization of the suspension and after 30 min rest. Finally, 100 μL of FBS were added into 900 μL of the previous suspension and the D_H_ was measured once again, after homogenization.

### 2.6. Small-angle X-Ray Scattering measurements

SAXS experiments on PEI/peGFP-C3 polyplexes were conducted using a XENOCS Xeuss 2.0 instrument, equipped with a GENIX-3D copper source, FOX-3D optics, and XENOCS motorized ‘anti-diffusion’ slits, delivering an 8 keV beam. The polyplex dispersions were placed into thin quartz capillaries (1.5 mm optical path, WJM-Glas/Müller GmbH, Germany), and data was collected over a *q* range of 0.004 to 0.05 Å⁻¹ for 3 hours. For these experiments, polyplexes were prepared using a peGFP-C3 of 0.34 mg/mL. Necessary corrections were applied, and the resulting data was analyzed using SASView software.

### 2.7. Atomic Force Microscopy

PEI/peGFP-C3 polyplexes were deposited on mica surfaces and imaged under ambient conditions at room temperature using a Dimension ICON AFM (Bruker, Billerica, USA). PeakForce mode was selected to carry on the imaging of PEI/peGFP-C3 polyplexes, allowing for precise control over the force exerted on the chains (setpoint) with SCANASYST-AIR probes. Tip radius of approximately 1–2 nm (Bruker, Billerica, MA, USA), and a scanning rate of 0.4 Hz were selected. Mica substrates were freshly cleaved before each experiment to ensure a smooth surface. PEI/peGFP-C3 polyplexes were prepared by solvent casting from a dilute stock solution (0.034 mg/mL) to reduce aggregation and promote the isolation of complexes on the mica surface for improved imaging. 4 µL of the suspension were applied to the freshly cleaved mica and allowed to dry under a nitrogen flow for a few minutes. The polyplexes were positively charged in solution at the selected formulation conditions, which facilitated strong electrostatic interactions with the negatively charged mica surface. The particle size distribution was assessed by measuring individual polyplexes from AFM micrographs using Image-J software.

### 2.8. HEK293T cell transfection

The *in vitro* studies were carried out using the HEK293T cell line [45] derived after stable expression of the SV40 large T antigen in the original immortalized HEK cells [46]. These cells exhibit rapid cell growth and are very permissive to transfection. HEK293T cells were cultured in DMEM supplemented with 10 % FBS, 2 mM L-Glutamine, and antibiotics (1% of stock solution, 15070-063, Gibco). Every week, the cells are detached using a trypsin solution (0.05%, Gibco) and are seeded at a density of 2 x 10^4^ cells/cm². The culture media are renewed every 2 days. For GFP expression analysis and cell viability assay (ATP relative content), HEK293T cells were seeded the day before transfection in 24- or 96-well plates at a density of 2 x 10^4^ cells/cm². For internalization tracking analysis, HEK293T cells were seeded the day before transfection in 48-well plates at the same density of 2 x 10^4^ cells/cm². The cell transfections were all carried out in triplicate. PEI/peGFP-C3 polyplexes were prepared with the four PEI samples at four selected charge ratios (R = [N^+^]/[P^−^] = 3, 5, 10 and 15), using the “one-shot and vortex” technique. Prior to transfection, polyplexes were all diluted in ultra-pure water up to a plasmid concentration of 0.022 mg/mL to have a constant amount of DNA among all the wells (2 μg). Then, a volume of 10 μL of PBS 10X was added to each diluted formulation. The culture medium from the wells was then removed and replaced with 900 μL of DMEM containing 10 % FBS and 100 μL of each formulation were added to each well. The plates were then incubated at 37 °C overnight. At 24 h after transfection, culture medium from the wells was discarded and replaced by regular supplemented DMEM supplemented medium.

### 2.9. Cytometry analysis and fluorescence microscopy

Cell monolayers were washed with PBS before observations by fluorescence microscopy using a Zeiss inverted microscope and the AxioVision software. Then, the cells were detached with trypsin-EDTA and resuspended in complete medium for flow cytometry analysis using a Becton Dickinson LSRFortessa™ X-20 (cytometry core facility of the Biology and Health Federative research structure Biosit, Rennes, France). Dot plots of forward scatter (FSC: x axis) and side scatter (SSC: y axis) allowed to gate the viable single cells (**Figure S.I. 1**). Untreated cells were used to determine autofluorescence, arbitrary set at ∼72 for HEK293T cells. The fluorescence emitted by GFP positive cells (at least 5,000 cells) was measured using the FITC-A channel and the fluorescence emitted cells containing cyanine 5-labelled-peGFP-C3 plasmid were detected using the APC-A channel. To analyze transfection efficiency, two parameters were quantified: the percentage of GFP or cyanine 5 positive cells and the mean of fluorescence (fluorescence intensity) reflecting the relative GFP expression levels in the cell population or the relative accumulation of cyanine 5-tagged plasmid. Flow cytometry data were analyzed using FlowLogic software (7.2.1 version, Inivai Technologies, Mentone Victoria, Australia).

### 2.10. ATP analysis

Cell viability was assessed by measuring the relative intracellular ATP content using CellTiter-Glo Luminescent Cell viability assay (Promega). At 24 and/or 48 hours after transfection, cells were incubated with the CellTiter-Glo reagent for 10 minutes (min). Lysates were transferred to an opaque multi-well plate and luminescent signals were quantified at 540 nm using the Polarstar Omega microplate reader (BMG Labtech). Cell viabilities in treated cells were expressed as the percentage of the luminescent values obtained in untreated cells, which was arbitrary set as 100%.

### 2.12. Laser Scanning Confocal Microscopy (LSCM)

Laser Scanning Confocal Microscopy Images were acquired on a Leica SP8 inverted microscope equipped with a 63/1.4 (oil immersion) objective in fluorescence mode. For these experiments, cells were plated in Lab-Tek™ II Chamber Slide™ (8 wells, reference 15534 from ThermoScientific™ Nunc™) at the same cell density as for other assays. At 24 or 48 hours, cells were fixed with 4% paraformaldehyde (16% formic aldehyde methanol free, reference 28908, ThermoScientific™ Pierce™) buffered with PBS (1/4 dilution vol/vol) for 15 min at 4°C. Paraformaldehyde was discarded and cell monolayers were rinsed twice with 1 X PBS. Then, cell nuclei were stained for 10 min using 40 mM Hoechst 33342 diluted in PBS (20 mM stock solution, reference 62249, ThermoScientific™) at room temperature. The Hoechst staining solution was discarded, the wells of the chamber slide device were removed, one drop of mounting solution (Vectashield antifade mounting medium, reference H-100à-10, Vector Laboratories) was dropped on the cell monolayers attached to the glass slide and the slide was fixed using a coverslip with nail polish prior to the observation of cells in confocal microscopy.

The laser outputs are controlled via the Acousto-Optical Tunable Filter (AOTF) and photomultiplicators (PMT) as follows: Hoechst 33342 was excited with a laser diode at 405 nm (17-22%) and measured with an emission setting at 410-443 nm; Cy5 was excited with an Helium-Neon laser at 633 nm (6-10%) while the emission was collected in the 639-676 nm range; GFP was excited with an Argon laser at 488 nm (15-20%) while the emission was collected in the 495-544 nm range. Transmission images were collected on the transmission photomultiplier tube (T-PMT) using Helium-Neon laser at 633 nm in transmission mode. Images were collected using the confocal microscope in sequential image recording to avoid the overlapped emission with a line average of 1 and a format of 512*512 pixels. Photomutliplicators gain and offset configurations were set up on control cells so as to correct images from green auto-fluorescence signal. Fluorescence confocal acquisitions and 3D reconstruction images were processed using ImageJ software.

### 2.13. Statistics

All quantitative data were expressed as mean ± standard deviation (SD). Experiments (n= independent experiments) were performed at least three times. Statistics were done using GraphPad Prism version 5.0 (GraphPad Software, USA). Differences between two groups were analyzed using two-tailed Mann-Whitney *U* test. A non-parametric Kruskal-Wallis test with Dunns’ post-test was used to compare means of more than two groups. Significant differences are presented as * p < 0.05, ** p < 0.01, *** p < 0.001, ^ns^ not significant.

## 3. Results and Discussion

### 3.1. PEI/peGFP-C3 polyplexes complexation and charge stoichiometry

The formation of PEI/peGFP-C3 polyplexes using the four PEI samples (*b*-PEI_0.8_, *l*-PEI_20_, *b*-PEI_25_ and *b*-PEI_60_) was monitored through ζ-potential measurements. The PEI solutions were systematically adjusted to pH 7. 4 before use. **Figure 1** shows the ζ-potential evolution as a function of charge ratio ([N^+^]/[P^−^]) during the progressive addition of small volumes of each PEI to a peGFP-C3 solution under continuous stirring. For DNA, since phosphate groups are fully ionized at neutral pH, [P^−^] is assumed to be equal to the molar concentration in nucleotides. For PEI, potentiometric titrations performed on all samples allowed determining the degree of protonation (σ) as function of pH. The values of σ at pH 7.4 are 0.47 for *b*-PEI_0.8_, 0.40 for *l*-PEI_20_, 0.30 for *b*-PEI_25_ and 0.33 for *b*-PEI_60_ (**Figure S.I. 2**).

**Figure 1.-.**
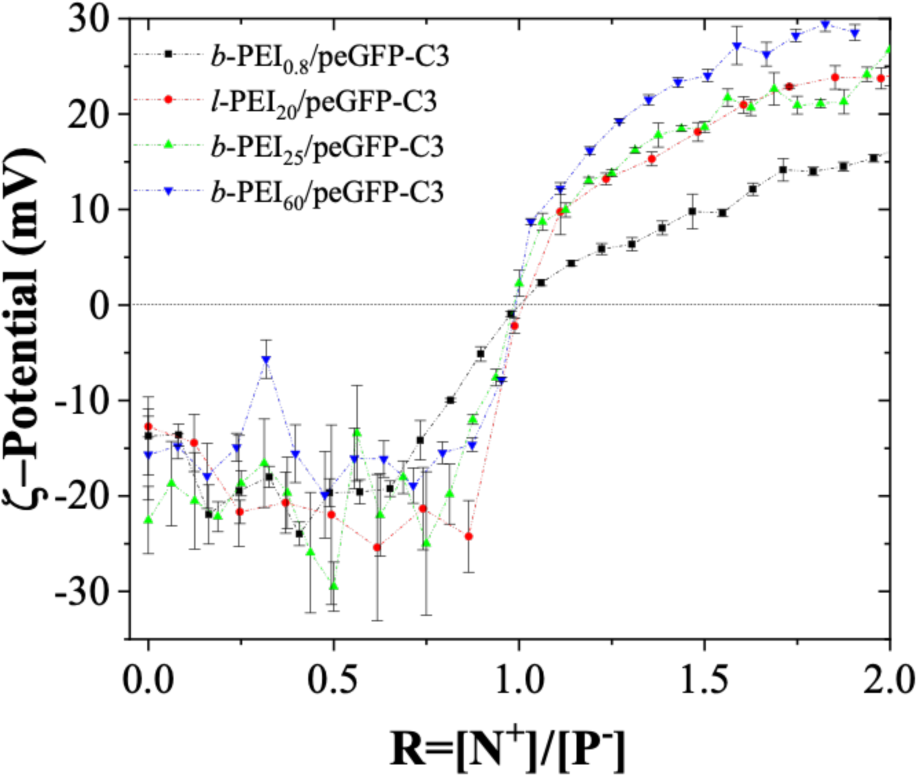
ζ-potential evolution as a function of charge ratio (R=[N^+^]/[P^−^]) during PEI/peGFP-C3 polyplexes formation using the slow dropwise mixing method and the four different PEI (Mw: 0.8, 20, 25 and 60 kg/mol). C_peGFP-C3_ = 0.005 mg/mL in water, C*_b_*_-PEI0.8_ = 0.8925 mg/mL, C*_l_*_-PEI20_ = 0.8721 mg/mL, C*_b_*_-PEI25_ = 0.9295 mg/mL and C*_b_*_-PEI60_ = 0.9700 mg/mL in water (pH 7.4). ζ-potential measurements were performed in triplicate and each titration was carried on by duplicate.

A plateau with negative ζ-potential values around −20 mV was observed for all PEI/peGFP-C3 polyplexes at a charge ratio R lower than 1. This plateau is related to the formation of partially complexed PEI/peGFP-C3 polyplexes with excess DNA. This is in good agreement with several reports in the literature for chitosan/DNA and cationic ELPs/DNA polyplexes [18,47–49]. Full complexation of peGFP-C3 negative charges, corresponding to a ζ-potential of 0 mV (isoelectric point) was achieved at R=[N^+^]/[P^−^]=1 for all PEIs. This indicates that the complexation between the positive charges of PEI and negative charges of DNA is stoichiometric. Thus, ζ-potential results clearly indicate that the complexation mechanism of PEI with peGFP-C3 is independent of PEI Mw and structure (linear vs. branched). However, ζ-potential values obtained in presence of an excess of PEI, *i.e.,* R > 1, increase from + 15 mV to around + 30 mV, with the lowest value corresponding to the smallest PEI Mw and increasing with Mw. The *l*-PEI_20_ and *b*-PEI_25_ exhibit similar ζ-potential, emphasizing that the Mw, rather than the structure, determines the surface charge of polyplexes. For R values higher than 1, PEI in excess is partially complexed at the surface of polyplexes, forming a corona of charged segments. The thickness of this layer directly correlates with the Mw of the PEI. Polyplexes with a larger corona of excess PEI are expected to be the most stable towards aggregation and plasmid release due to enhanced electrostatic repulsion. Our results are in good agreement with reports in the literature for PEI/DNA polyplexes prepared with PEI samples with Mw of 2, 5, and 25 kg/mol, where PEI 25 kg/mol was found to present the highest positively surface charged polyplexes with DNA [50].

The binding properties of peGFP-C3 with the different PEI samples were also studied using gel electrophoresis experiments, which were performed by analyzing the electrophoretic mobility of the plasmid DNA at different charge ratios R ([N^+^]/[P^−^]) on the agarose gel (**Figure 2**). **Figures 2a** to **d** present the gel electrophoresis results for *b*-PEI_0.8_, *l*-PEI_20_, *b*-PEI_25_ and *b*-PEI_60_/peGFP-C3 polyplexes, respectively, showing the effect of progressive compaction of peGFP-C3 at increasing concentrations of the respective PEI sample. It can also be observed the predominance of the supercoiled conformation in the initial peGFP-C3 plasmid, while during its complexation with *b*-PEI_0.8_ and *l*-PEI_20_, an increase on the intensity of the band corresponding to coiled conformation is observed. Our results show that PEI can effectively neutralize peGFP-C3 from a charge ratio of 1, which is in good agreement with the ζ-potential measurements previously discussed and presented in **Figure 1**. These results allowed determining the experimental stoichiometry in order to prepare PEI/peGFP-C3 polyplexes with a plasmid concentration of 0.034 mg/mL and R > 1, using the “one shot and vortex” rapid mixing method, for the stability studies and biological assays presented in the following sections of this paper.

**Figure 2.-.**
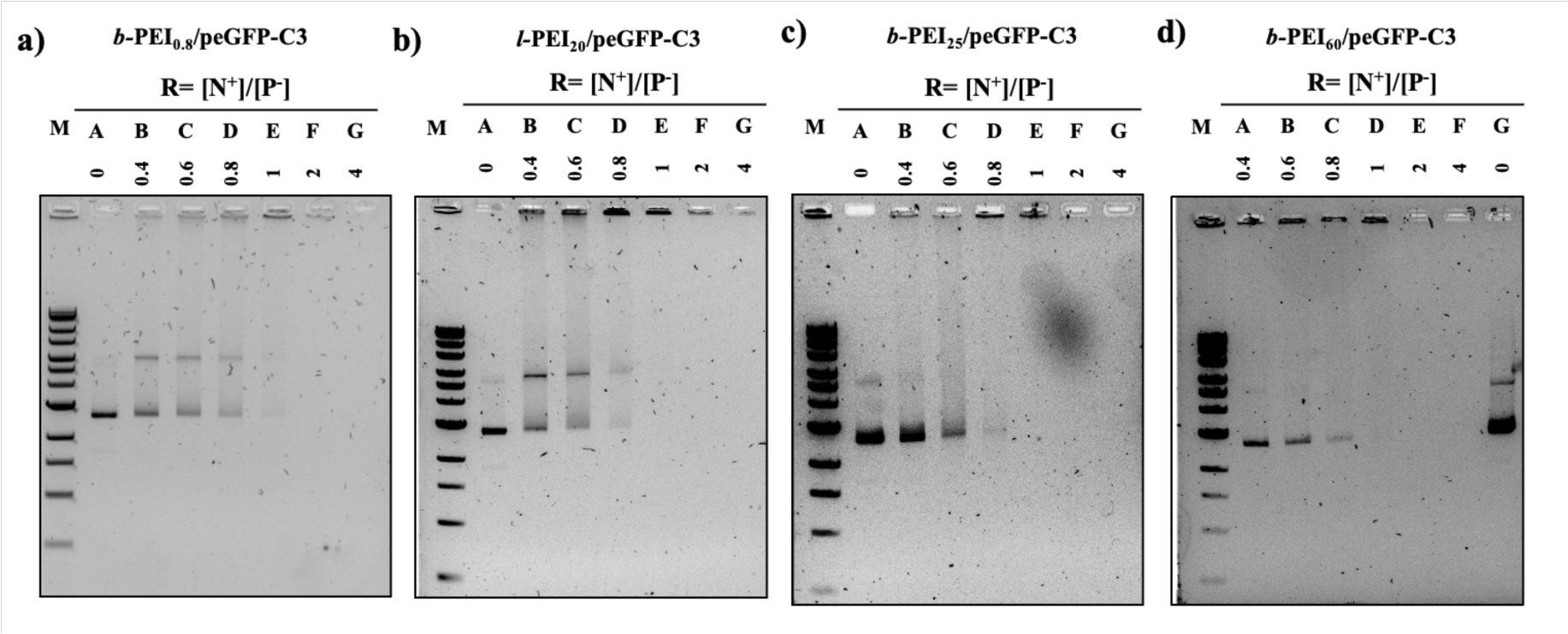
Electrophoresis gel assays showing the effect of charge ratio, [N^+^]/[P^−^], on the electrophoretic mobility of eGFP-C3 plasmid in presence of a) *b*-PEI_0.8_, b) *l*-PEI_20_, c) *b*-PEI_25_ and d) *b*-PEI_60_. M shows pDNA markers of different Mw (SmartLadder Mw-1700-10 Eurogentec). C_peGFP-C3_ = 0.005 mg/mL prepared in water. Mother solutions of PEI were prepared at the following concentrations: C*_b_*_-PEI0.8_ = 0.019 mg/mL, C*_l_*_-PEI20_ = 0.024 mg/mL, C*_b_*_-PEI25_ = 0.024 mg/mL and C*_b_*_-PEI60_ = 0.02 mg/mL in water (pH 7.4).

### 3.2. PEI/peGFP-C3 polyplexes characterization

Subsequent PEI/peGFP-C3 polyplexes were prepared in DI water by quickly adding the PEI solution into the plasmid solution with a micropipette, followed by an immediate vortex to mix both solutions during 10 seconds. AFM measurements were employed to image and analyze the morphology of dried polyplexes suspensions’ morphology of *b*-PEI_25_/peGFP-C3 polyplexes at a charge ratio of R = 4 (**Figures 3a** and **b**). **Figure 3a** shows the presence of particles with sizes around 50 nm and **Figure 3b** presents a zoom on two polyplexes, revealing the existence of a hierarchical complex structure, where small primary polyplexes formed during the initial phases of complexation aggregate to create bigger secondary polyplex particles. This morphology was also reported and described for lysozyme /poly(styrene sulfonate) (PSS) [51], PEI/poly(styrene sulfonate) (PSS) [52] and PSS/poly(diallyldimethylammonium chloride) (PDADMAC) polyplexes [53]. **Figure 3c** shows the ζ-potential values for PEI/peGFP-C3 polyplexes prepared with the four PEI samples at the charge ratios from 0.5 to 15. As expected, the ζ-potential values are negative for PEI/peGFP-C3 polyplexes prepared at R<1 due to the excess of peGFP-C3 negatively charged chains, while they are positive for polyplexes prepared at R values higher than the stoichiometry (R>1), due to the excess of positively charged PEI chains. The full complexation state of the plasmid is found at the highest positive ζ-potential values for each PEI sample. Furthermore, the ζ-potential values for PEI/peGFP-C3 polyplexes, with R > 1, were found to increase with the increment of the Mw of PEI: *b*-PEI_60_ > *b*-PEI_25_ > *b*-PEI_0.8_; in good agreement with the previous titrations and with reports in the literature [50]. The ζ-potential values for *l*-PEI_20_/peGFP-C3 and *b*-PEI_25_/peGFP-C3 polyplexes were found to be quite similar for both particles.

**Figure 3.-.**
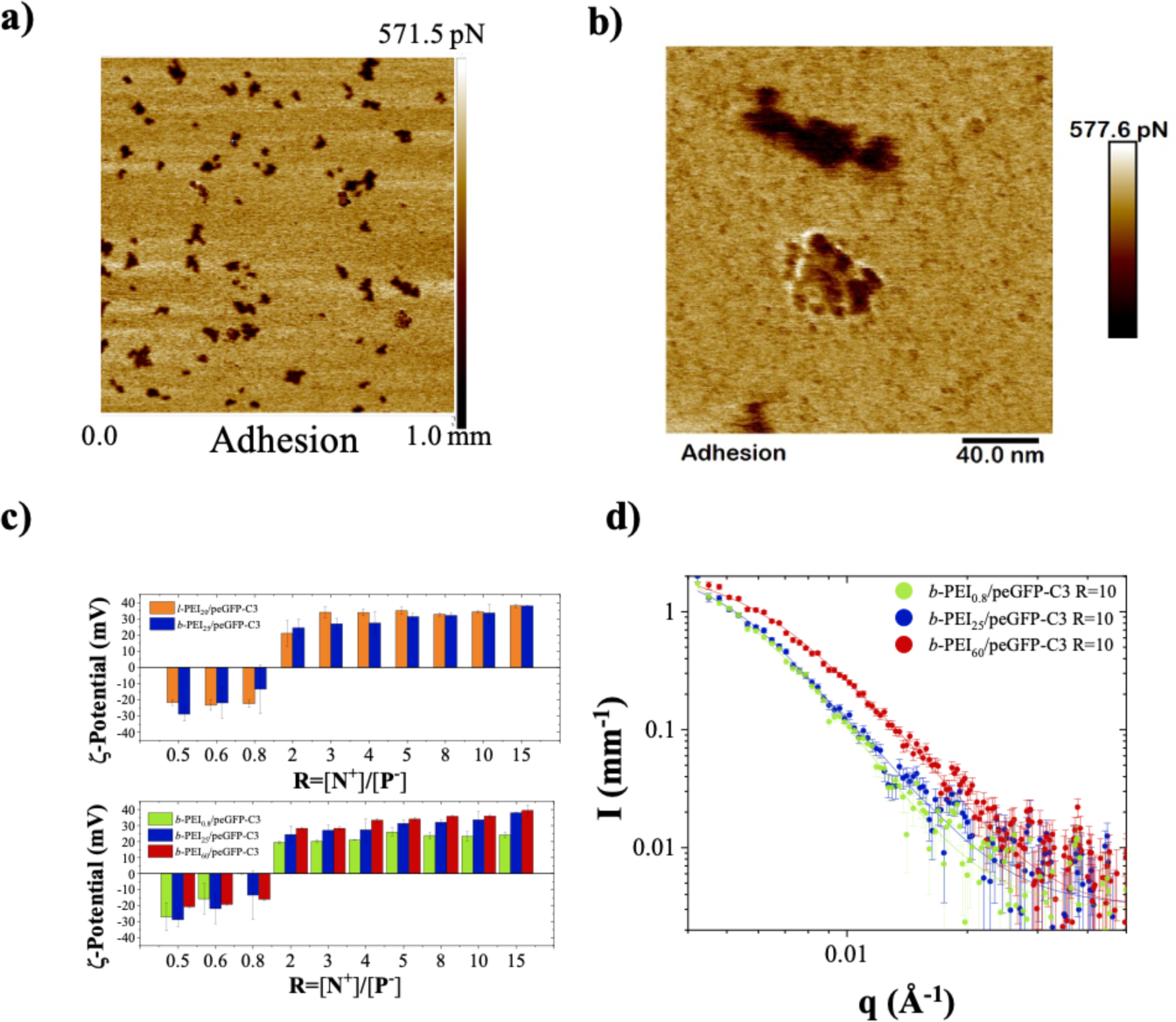
a) and b) AFM images for *b*-PEI_25_/peGFP-C3 polyplexes prepared at a charge ratio of [N^+^]/[P^−^]= 4. c) ζ-potential as a function of [N^+^]/[P^−^] ratio for PEI/peGFP-C3 polyplexes prepared with the different PEI samples: *b*-PEI_0.8_, *l*-PEI_20_, *b*-PEI_25_ and *b*-PEI_60_. d) SAXS intensity I(q) as a function of the wave vector *q* for PEI/peGFP-C3 polyplexes prepared with *b*-PEI_0.8_, *b*-PEI_25_ and *b*-PEI_60_ at R=10. For AFM and ζ-potential measurements: C_peGFP-C3_= 0.034 mg/mL in water, C*_b_*_-PEI0.8_ = 0.019 mg/mL, C*_l_*_-PEI20_ = 0.024 mg/mL, C*_b_*_-PEI25_ = 0.024 mg/mL and C*_b_*_-PEI60_ = 0.02 mg/mL in water (pH 7.4). For SAXS measurements: C_peGFP-C3_= 0.340 mg/mL in water. ζ-potential experiments were performed in triplicate. All polyplexes were prepared through rapid one-shot mixing of PEI to eGFP-C3 plasmid.

**Figure 3d** presents the background-subtracted SAXS data for PEI/peGFP-C3 polyplexes prepared with the branched PEI samples at a charge ratio of 10. Based on AFM images suggesting a hierarchical complex structure, where small primary polyplexes aggregate into larger secondary polyplex particles, the data were fitted using a polydisperse (hairy) sphere form factor [54]. This analysis provided information upon the radii of the primary polyplexes, which were determined to be 19.7 ± 8.9 nm for *b*-PEI_0.8_/peGFP-C3, 7.9 ± 5.5 nm for *b*-PEI_25_/ peGFP-C3 and 9.1 ± 5.4 nm for *b*-PEI_60_/peGFP-C3. The results indicate that denser polyplexes are formed with *b*-PEI_25_ and *b*-PEI_60_ compared to those with *b*-PEI_0.8_/peGFP-C3 polyplexes. Additionally, a slightly larger radius (10.5 ± 6.3 nm) was observed for *b*-PEI_25_/peGFP-C3 polyplexes prepared at R=0.6 compared to those at R=10 (7.9 ± 5.5 nm), suggesting that a substantial excess of b *b*-PEI_25_ is required to effectively compact the plasmid into small, dense particles.

**Figure 4** depicts the size distribution of different polyplexes prepared with the four PEI samples at four charge ratios (3, 5, 10 and 15) analyzed with dynamic light scattering (DLS). Whatever the PEI used, polyplexes present sizes below 100 nm and demonstrate a relatively small size distribution, with PDI lower than 0.3. These polyplexes are expected to exhibit a morphology characterized by a «core» predominantly made up of the peGFP-C3 complexed with either branched or linear PEI, and a «corona» that contains an excess of partially complexed PEI, where free protonated amines contribute to the polyplexes positive charge. As the R value increases, polyplexes become progressively more compact and present lower particles sizes, in good agreement with reports in the literature for other systems such as chitosan/DNA and ELPs/DNA polyplexes [18,47,48], suggesting that a significant excess of PEI is necessary to produce small and dense particles for an effective condensation of peGFP-C3. Furthermore, among polyplexes prepared with the branched PEI samples, the higher particle sizes were obtained for *b*-PEI_60_/peGFP-C3 polyplexes. This trend was also observed by other authors working with chitosan/DNA polyplexes [55,56]. They suggested that at high Mw, limitations of the polycation chain upon complexation become more important, giving rise to a different complexation behavior for high Mw polycations compared to low ones. Finally, particle sizes obtained for *l*-PEI_20_/peGFP-C3 polyplexes are slightly larger than those of *b*-PEI_25_/peGFP-C3 polyplexes. This suggests that particles prepared with linear PEI are looser and presumably less stable than those formulated with branched PEI.

**Figure 4.-.**
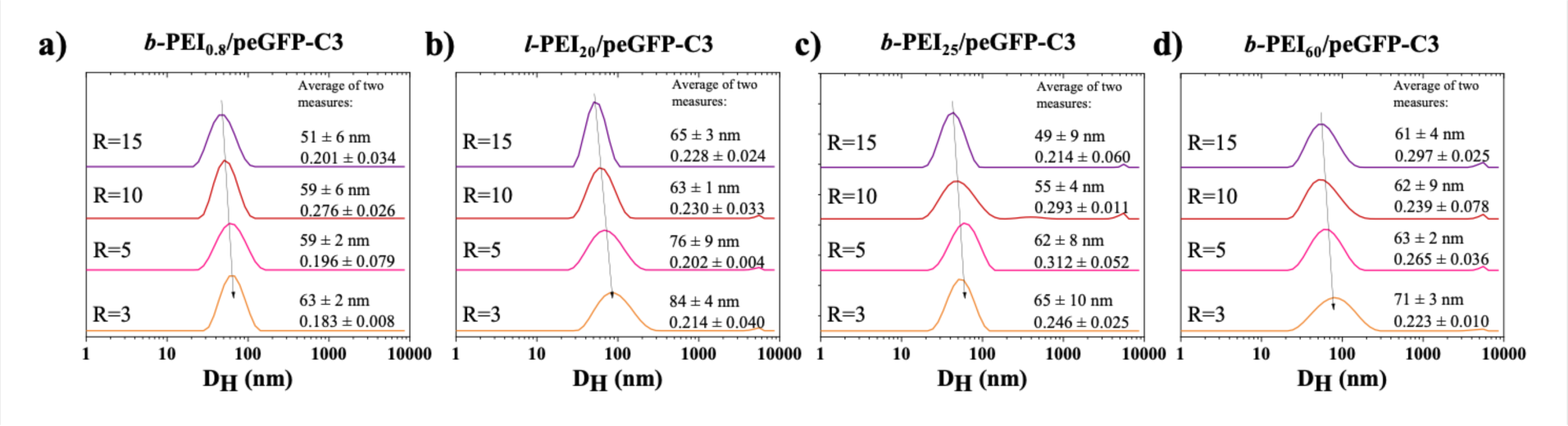
Intensity-averaged size (hydrodynamic diameter) distributions obtained by DLS for PEI/peGFP-C3 polyplexes prepared at different charge ratios ([N^+^]/[P^−^]= 3, 5, 10 and 15) using the following PEI samples: a) *b*-PEI_0.8_, b) *l*-PEI_20_, c) *b*-PEI_25_ and d) *b*-PEI_60_. C_peGFP-C3_ = 0.034 mg/mL in water, C*_b_*_-PEI0.8_ = 0.8925 mg/mL, C*_l_*_-PEI20_ = 0.8721 mg/mL, C*_b_*_-PEI25_ = 0.9295 mg/mL and C*_b_*_-PEI60_ = 0.9254 mg/mL in water (pH 7.4). Temperature measurement: 25 °C. DLS measurements were performed in duplicate. PEI/peGFP-C3 polyplexes were prepared by using the rapid one-shot and vortex method.

### 3.3. PEI/peGFP-C3 polyplexes stability in biological media

Polyplexes stability was studied in terms of size evolution in the presence of biological media (PBS, DMEM and DMEM supplemented with 10% of fetal bovine serum) by DLS (**Figures 5**), as well as with gel electrophoresis to determine if peGFP-C3 plasmid was burst released at those conditions even before being internalized (**Figure S.I. 3**).

**Figure 5.-.**
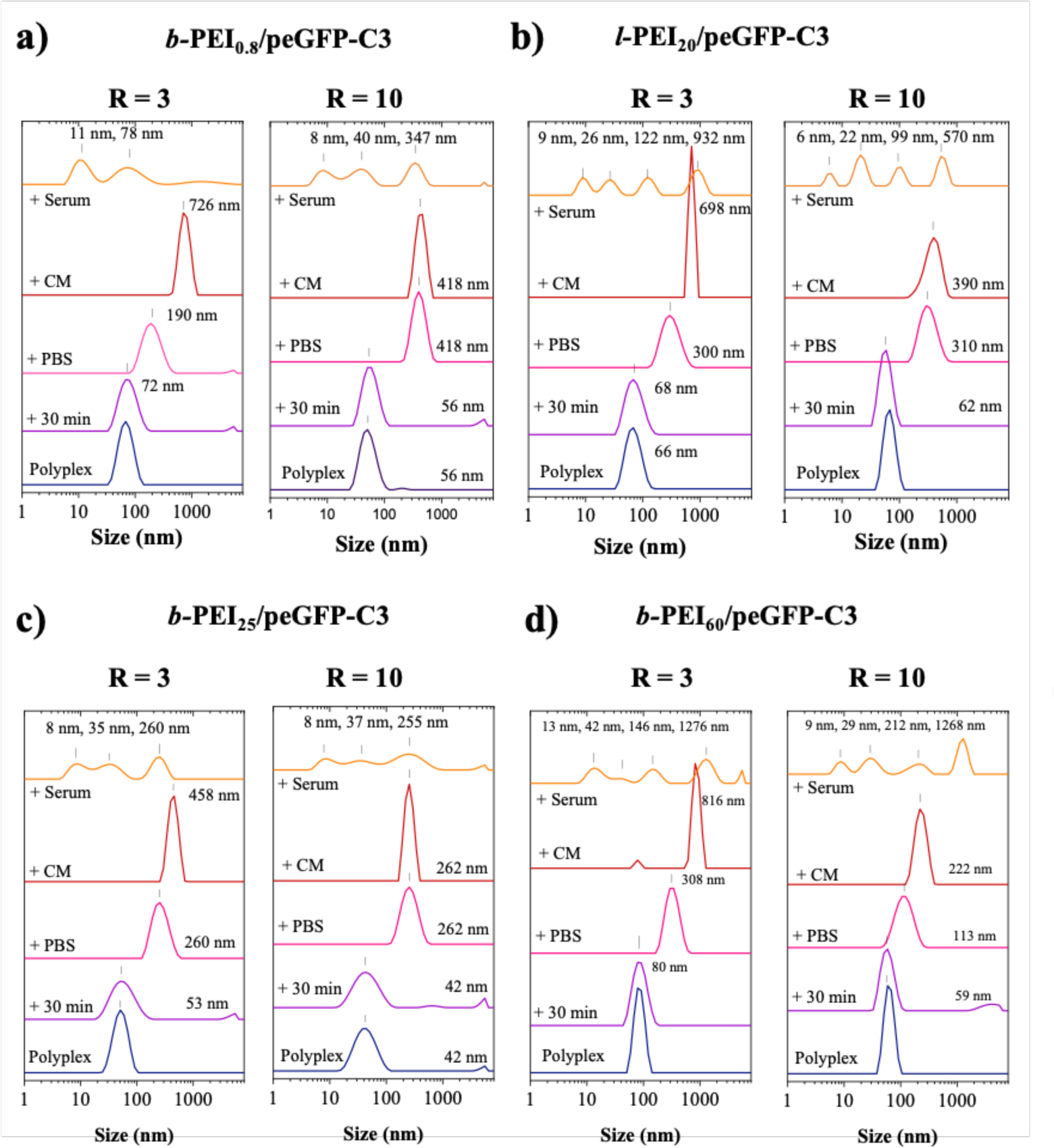
Intensity-averaged size distribution obtained by DLS for PEI/peGFP-C3 polyplexes prepared at two different charge ratios ([N^+^]/[P^−^]= 3 and 10, using the following PEI samples: a) *b*-PEI_0.8_, b) *l*-PEI_20_, c) *b*-PEI_25_ and d) *b*-PEI_60_, in presence of PBS, PBS with culture media (CM) and PBS, culture media and serum. Polyplexes hydrodynamic diameter was determined just after formulation (t=0) and at t=30 min. Temperature measurement: 25 °C. C_peGFP-C3_ = 0.034 mg/mL in water, C*_b_*_-PEI0.8_ = 0.8925 mg/mL, C*_l_*_-PEI20_ = 0.8721 mg/mL, C*_b_*_-PEI25_ = 0.9295 mg/mL and C*_b_*_-PEI60_ = 0.9254 mg/mL in water (pH 7.4). Dulbeccos Modified Eagle Medium was used as the culture medium and fetal bovine serum (FBS) at 10% was used as the serum. Samples with PBS were prepared with 100 µL of initial polyplex, 100 µL of PBS 10x and 800 µL of ultrapure water. Samples with CM were prepared with 500 µL of the previous suspension containing PBS and mixed with 500 µL of culture medium. Samples with CM and serum were prepared with 900 µL of precedent solution and 100 µL of serum. DLS measurements were performed in duplicate.

**Figure 5a** shows the intensity-averaged size distribution evolution of *b*-PEI_0.8_/peGFP-C3 polyplexes prepared at R= 3 and 10, just after polyelectrolytes rapid mixing formulation technique, 30 min after preparation, and then in presence of PBS, PBS and CM and finally, with serum. It is well known that the addition of salt diminishes both the attractive interactions between the oppositely charged polyelectrolytes and the repulsive forces among the polyplexes, possibly leading to the dissociation of the polyplexes and/or their aggregation [57–59]. We can observe that polyplexes D_H_ is quite stable after 30 min of preparation, even after 7 weeks (data not shown). Then, in presence of PBS, D_H_ increase up to 5 times, probably due to the decrease of electrostatic attractive interactions between peGFP-C3 and *b*-PEI_0.8_, or by polyplexes aggregation, resulting from the presence of salts in PBS (ionic strength 162.7 mM). D_H_ of *b*-PEI_0.8_/peGFP-C3 polyplexes prepared at a R of 3 increases again after the addition of CM, revealing a low stability for low Mw PEI-based polyplexes at this charge ratio. However, for R equal to 10, sizes remain the same, showing that at this charge ratio, polyplexes remain stable in presence of CM. Finally, after the addition of serum, different sized populations are observed, most likely due to the presence of proteins in the serum, making the DLS analysis difficult and without possibility to conclude about polyplexes stability. In fact, the intensity-averaged size distribution evolution of serum alone already presents multiple sized populations (**Figure S.I. 4**).

Furthermore, gel electrophoresis assays of *b*-PEI_0.8_/peGFP-C3 polyplexes prepared at the charge ratios of 3 and 10 studied in presence of the different biological media revealed that some plasmid was released from polyplexes prepared at R=3 subjected to the presence of PBS and CM (**Figure S.I. 3a**). It is worth mentioning that a band related to remaining DNA in the serum is present in the lines of polyplexes with PBS, CM and serum, as well as in the line of the control containing PBS and CM, in the presence of serum demonstrating the presence of free genomic DNA in the serum. After analyzing the stability of *b*-PEI_0.8_/peGFP-C3 polyplexes, it is possible to conclude that low Mw PEI (*b*-PEI_0.8_) does not provide enough stability to polyplexes in presence of biological media, giving sizes up to 726 nm in PBS and CM at charge ratio of 3. This emphasizes the importance of the Mw and amine content of the polymeric gene carrier, and particle size under physiological conditions, impacting certainly the polyplexes at the early stages of cell internalization [23,36].

**Table 1** summarizes the values of the hydrodynamic diameter for PEI/peGFP-C3 polyplexes prepared with the four PEI samples at the charge ratios of 3, 5, 10 and 15, and when diluted with PBS, CM and serum.

**Table 1.**
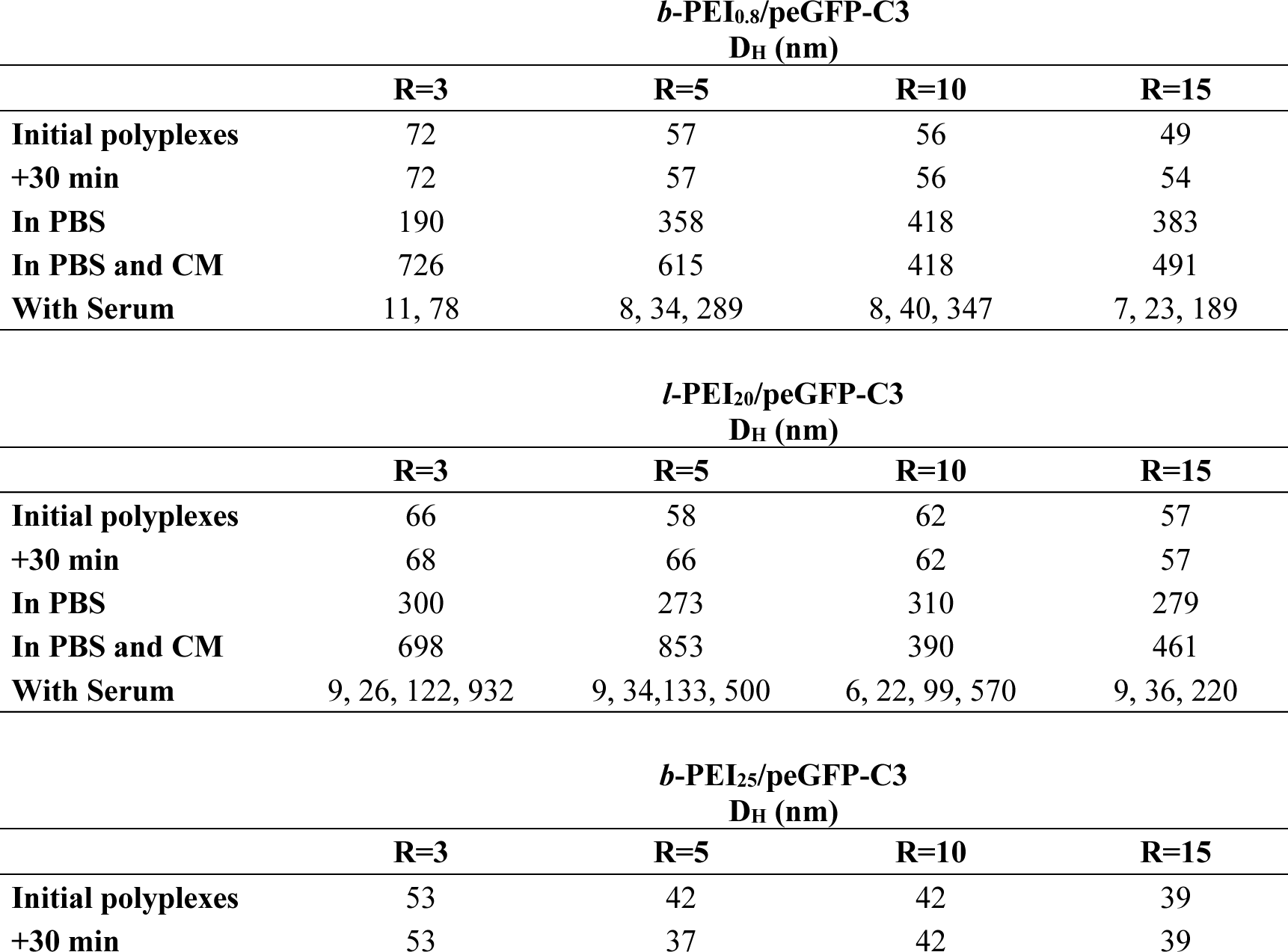
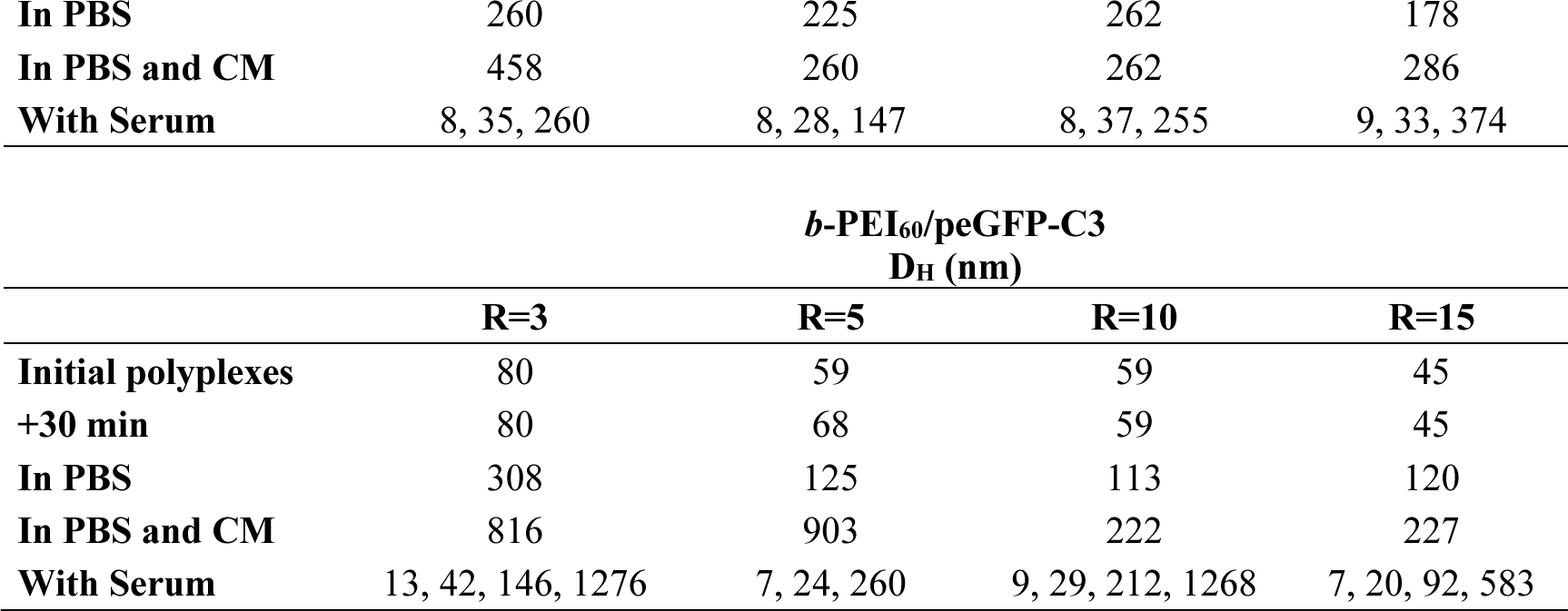
Evolution of hydrodynamic diameter (D_H_) of *b*-PEI_0.8_, *l*-PEI_20_, *b*-PEI_25_ and *b*-PEI_60_/peGFP-C3 polyplexes measured immediately after preparation, then, 30 min after preparation and in presence of PBS, CM and serum.

**Figure 5b, 5c** and **5d** show the same study carried on for *l*-PEI_20_/peGFP-C3, *b*-PEI_25_/peGFP-C3 and *b*-PEI_60_/peGFP-C3 polyplexes prepared at R= 3 and 10, in presence of PBS, PBS with CM, and PBS with CM and serum. A similar trend is observed for all polyplexes, this is, the particle size remains the same 30 min after polyplexes preparation and it increases with the addition of PBS and CM. However, it was found that stability in terms of size is highly dependent on polyplexes charge ratio and PEI’s Mw. Thus, polyplexes prepared at a charge ratio of 10 present a better stability than those prepared at the charge ratios of 3 and 5. Furthermore, branched PEI samples lead to the formation of polyplexes owning a greater stability than those prepared with the linear PEI sample, since particle sizes obtained after the addition of PBS and CM are larger for polyplexes prepared with *l*-PEI_20_. The branched PEI with the highest Mw (*b*-PEI_60_) likely promotes the formation of stable polyplexes, presenting a small size increase for the charge ratios of 10 and 15. From gel electrophoresis assays of *l*-PEI_20_/peGFP-C3, *b*-PEI_25_/peGFP-C3 and *b*-PEI_60_/peGFP-C3 polyplexes prepared at the charge ratios of 3 and 10 in presence of PBS, CM and serum, it was possible to observe that even if particle sizes increase, no plasmid was released from the polyplex (**Figure S.I. 3 a to d**), except for *b-*PEI_0.8_/peGFP-C3. Our results reveal that the presence of salt impacts the physicochemical properties of PEI/peGFP-C3 polyplexes, which is in good agreement with reports in the literature [65]. In general, polyplexes diluted with saline media show a lower stability and larger particle sizes than polyplexes prepared in water. However, this phenomenon can be tuned with the variation of the Mw of PEI, as well as the charge ratio of polyplexes.

### 3.4. Transfection efficiencies in HEK293T cells using PEI/peGFP-C3 polyplexes

Furthermore, we compared the transfection efficiencies in HEK293T cells obtained using the PEI/peGFP-C3 polyplexes prepared with the four PEI samples (*b*-PEI_0.8_, *l*-PEI_20_, *b*-PEI_25_ and *b*-PEI_60_) at four charge ratios (R = [N^+^]/[P^−^] = 3, 5, 10 and 15) and the lipofectamine 3000/peGFP-C3 lipoplexes being this transfection reagent widely used as a positive control (**Figure 6, Figure S.I. 5**).

**Figure 6.-.**
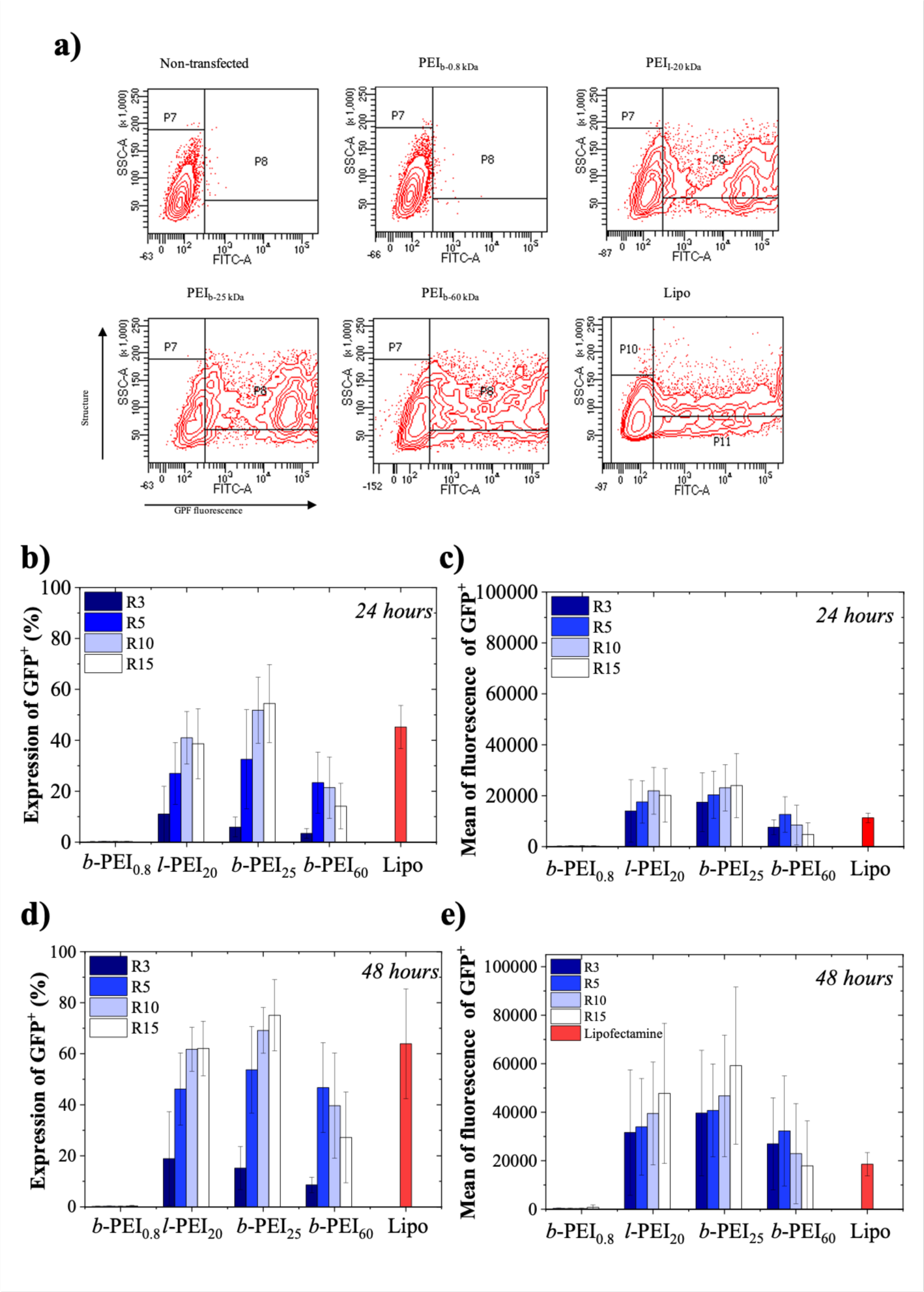
Transfection efficiency of HEK293T cells using *b*-PEI_0.8_, *l*-PEI_20_, *b*-PEI_25_ and *b*-PEI_60_/ peGFP-C3 polyplexes prepared at four different charge ratios ([N^+^]/[P^−^]= 3, 5, 10 and 15), and lipofectamine 3000/peGFP-C3 lipoplexes. Transfection efficiencies were evaluated after 24 and 48 hours post-transfection by flow cytometry (**a**) to determine both percentages of GFP positive cells (**b**, **d**) and means of fluorescence (**c**, **e)**. These data are the results of 5 independent experiments (n=5).

Our flow cytometry settings (**Figure S.I. 1**) allowed to quantify both percentages of GFP positive (GFP^+^) cells and the means of fluorescence in the GFP^+^ cell population, which reflects the relative level of GFP expression in these cells (**Figure 6a**). From these raw data of flow cytometry (**Figure 6a**), we drew histograms showing both parameters in HEK293T cells at 24 (**Figure 6b** and **6c)** and 48 h after transfection (**Figure 6d** and **6e**). Despite quantitative variations between the independent experiments performed, our results demonstrated that *l*-PEI_20_, *b*-PEI_25_, *b*-PEI_60_ and lipofectamine/peGFP-C3 complexes allowed efficient transfections after 24 h, which further increased at 48 h while no transfected cells were detected using *b*-PEI_0.8_/peGFP-C3 polyplexes. Percentages of GFP^+^ cells increased with higher R=[N^+^]/[P^−^] to reach 60 %, 70 % and 45 % of positive cells at 48 h with *l*-PEI_20_, *b*-PEI_25_, *b*-PEI_60_/peGFP-C3 polyplexes, respectively, at R=[N^+^]/[P^−^]= 10 and 15 while lipofectamine/peGFP-C3 lipoplexes allowed ∼60 % GFP^+^ cells (**Figure 6b** and **6d)**. The transfection efficiencies at R=[N^+^]/[P^−^]=3 were much lower and below 20 % for all polyplexes.

The means of fluorescence showed a nearly 3-fold increase between 24 and 48 h, but we found, however less differences in this parameter between the *l*-PEI_20_, *b*-PEI_25_, *b*-PEI_60_/peGFP-C3 polyplexes (**Figure 6c** and **6e)** strongly suggesting that the relative expressions of GFP in cells transfected with these different polyplexes were quite similar although the percentages of GFP^+^ cells significantly varied between PEI/peGFP-C3 polyplexes.

Despite high percentages of GFP^+^ cells, the means of fluorescence were slightly lower with lipofectamine/peGFP-C3 lipoplexes (**Figure 6c, e).** While transfection with *b*-PEI_0.8_/peGFP-C3 polyplexes had no significant effect on cell numbers, transfection with *l*-PEI_20_, *b*-PEI_25_ and *b*-PEI_60_/peGFP-C3 polyplexes reduced the cell viability in a concentration dependent manner (at high R = [N^+^]/[P^−^] = 10 and 15) at 24 hours after incubation (**Figure S.I. 6**). This decreased number of cells compared to control culture condition may be due in part to cell death but also to a cytostatic effect since after discarding the transfection media at 24 hours, the decrease in cell number was no longer observed at 48 hours.

### 3.5. Correlation between the internalization of PEI/peGFP-C3 polyplexes and GFP expression

In order to determine whether the differences in transfection efficiencies observed with the PEI/peGFP-C3 polyplexes and lipofectamine/peGFP-C3 lipoplexes resulted from different levels in intracellular uptake of these complexes, we next used cyanine 5-labeled peGFP-C3 plasmid (Cy5-peGFP-C3) complexed with PEI and lipofectamine to quantify by flow cytometry the accumulation of plasmid in HEK293T cells at different times after transfection (**Figure 7a**). First, we demonstrated that the cyanine 5 labeling of the peGFP-C3 plasmid resulted in a strong fluorescence signal in HEK293T cells that had internalized the PEI and Lipofectamine complexes. Unexpectedly, we observed that that nearly 50% of the cells had internalized *b*-PEI_0.8_, *l*-PEI_20_, *b*-PEI_25_ and *b*-PEI_60_ Cy5-peGFP-C3 polyplexes and lipofectamine 3000/ Cy5-peGFP-C3 lipoplexes as early as 1 hour after the incubation (**Figure 7a**). In contrast, no cyanine 5 positive (Cy5^+^) cells were detected when *b*-PEI_0.8_/Cy5-peGFP-C3 polyplexes were used. After 2 hours of incubation, nearly 50 % of the cells became Cy5^+^ with all the PEI and lipofectamine 3000/Cy5-peGFP-C3 complexes including with *b*-PEI_0.8_/Cy5-peGFP-C3 polyplexes. Then, at 8 hours of incubation, more than 95 % of the cells were Cy5^+^ with all the polyplexes. At 24 hours, the Cy5^+^ fluorescence signal increased in the cells transfected with the *l*-PEI_20_, *b*-PEI_25_ and *b*-PEI_60_/ and lipofectamine 3000/Cy 5-peGFP-C3 polyplexes but not for those prepared with *b*-PEI_0.8_. At 48 hours, the signal of fluorescence remained very high but some of the Cy5^+^ cells exhibited a weaker fluorescence compared to the histograms obtained at 24 hours of incubation, suggesting that a fraction of the internalized plasmid was degraded and/or diluted when HEK293T cells proliferates resulting in a reduced mean of Cy5^+^ fluorescence.

**Figure 7.-.**
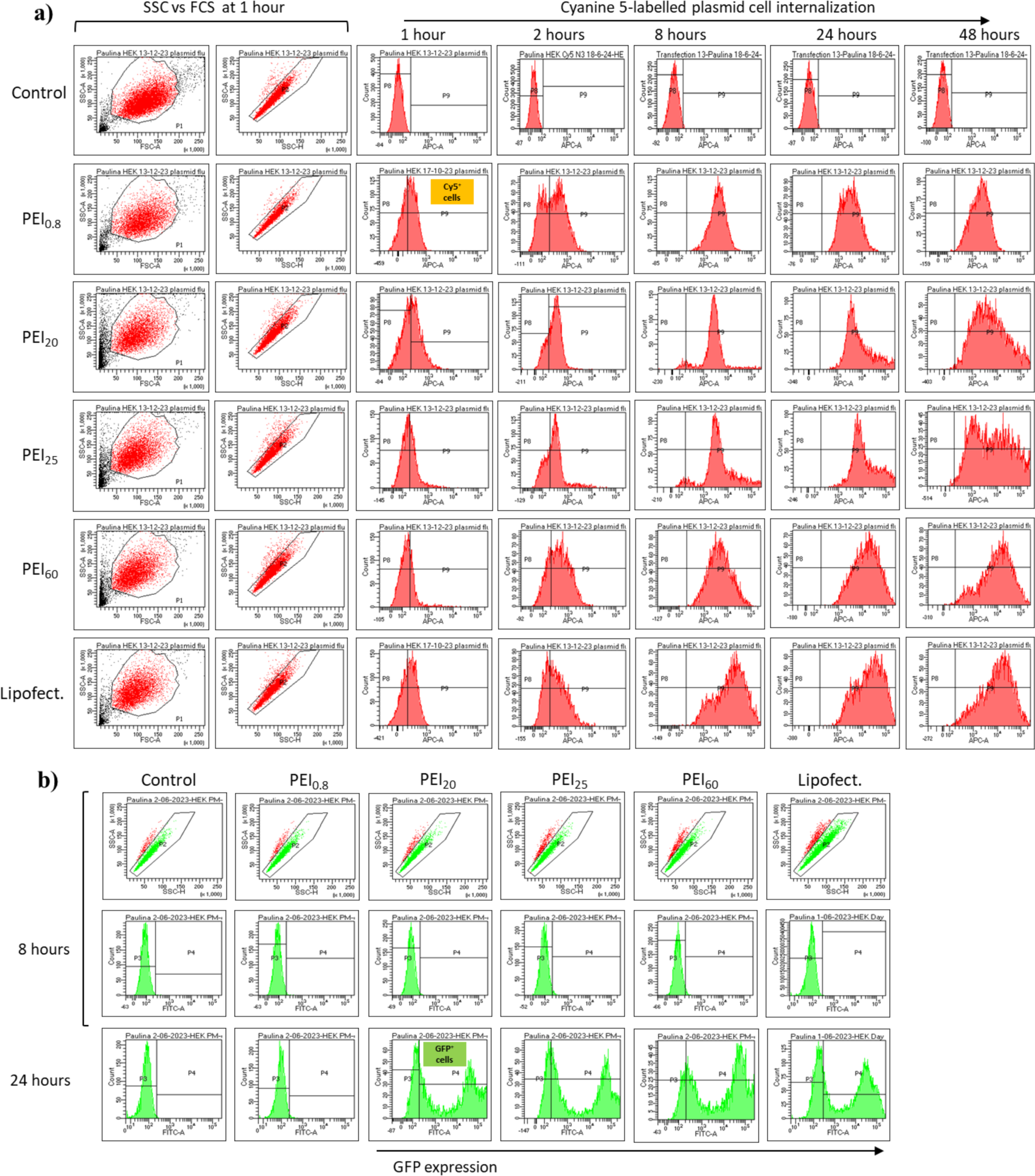
Flow cytometry analysis of HEK293T cells transfected with *b*-PEI_0.8_, *l*-PEI_20_, *b*-PEI_25_ and *b*-PEI_60_)/cyanine 5-peGFP-C3 polyplexes and lipofectamine/cyanine 5-peGFP-C3 lipoplexes (**a**) and PEI/ and lipofectamine/peGFP-C3 encoding GFP (**b**) at various time points after setting up the transfection procedure. Dot plots FSC-A vs SSC-A and SSC-H vs SSC-A represent the gates used to select viable cells and single cells, respectively. Histograms APC-A/counts and FITC-A/counts display the cells acquired to detect PEI or lipofectamine/cyanine 5-peGFP-C3 complexes and GFP^+^ cells, respectively.

It is also important to note that the fluorescence histograms at 24 and 48 hours characterizing the Cy5^+^ HEK293T cells incubated in presence of *l*-PEI_20_, *b*-PEI_25_ and *b*-PEI_60_/peGFP-C3 polyplexes and lipofectamine 3000/Cy5-peGFP-C3 were very wide with fluorescence intensity differences ranging from 2 to 3 log between cells. These data demonstrated that the intracellular amounts of plasmid varied considerably between the cells and greatly evolved during the incubation time with polyplexes and lipoplexes and after the renewal of the culture medium and the removal of the excess of polyplexes at 24 hours.

Concomitantly to these experiments studying the internalisation of Cy5-labeled peGFP-C3 complexed to PEI and Lipofectamine 3000, we also investigated the onset of the GFP expression (**Figure 7b**). Interestingly, we found that GFP was not expressed in HEK293T cells during the first 8 hours of incubation with polyplexes and lipoplexes despite the fact that most cells had internalized these complexes. The expression occurred between 8 and 24 hours after incubation but the percentages of GFP^+^ cells at 24 (**Figure 6b**) and 48 (**Figure 6c**) hours remained lower than the percentages of Cy5^+^ cells (**Figure 7a**) for HEK293T cells transfected with *l*-PEI_20_, *b*-PEI_25_ and *b*-PEI_60_/ Cy5-peGFP-C3 polyplexes and lipofectamine 3000/Cy5-peGFP-C3 lipoplexes. For the *b*-PEI_0.8_/peGFP-C3 polyplexes all cells were Cy5^+^ (**Figure 7a**) at 24 and 48 h but no GFP^+^ cells were detected (**Figure 6b, d**).

From the flow cytometry data (**Figure 7a**), we plotted the percentages and/or the means of fluorescence in both Cy5^+^ and GFP^+^ cells (**Figure 8**). We first drew the histograms presenting the percentages (**Figure 8a**) and means of fluorescence (**Figure 8b**) of Cy5^+^ and GFP^+^ cells at different time points after incubation with *b*-PEI_25_/(Cy5)-peGFP-C3 polyplexes at R=[N^+^]/[P^−^]=5, which gave high transfection efficiencies (**Figure 6**). As previously mentioned, 50 ± 10 % of cells were Cy5^+^ as early as 1 h after incubation with this polyplex and all cells became positive at 24 h while 25 % and 50 % of GFP^+^ cells were detected at 24 and 48 h, respectively (**Figure 8a**). Similarly, the means of Cy5^+^ progressively increased between 1 and 24 h of incubation, then slightly decreased at 48 h (**Figure 8b**). In order to evidence the quantitative differences in the accumulation of Cy5-peGFP-C3, we next represented the Cy5^+^ fluorescence means at 24 h of HEK293T cells transfected with *b-*PEI_25_/(Cy5)-peGFP-C3 polyplexes and lipofectamine/(Cy5)-peGFP-C3 lipoplexes from 5 independent experiments (**Figure 8c**). We evidenced large differences in the amounts of Cy5-peGFP-C3 internalized following incubation with PEI polyplexes and lipofectamine lipoplexes. The transfection using *b*-PEI_0.8_/Cy5-peGFP-C3 polyplexes gave the lowest mean of fluorescence, then, increasing values were obtained for *b*-PEI_25_, *l*-PEI_20_, *b*-PEI_60_, and lipofectamine 3000/Cy5-peGFP-C3 complexes. We next correlated these Cy5^+^ fluorescence means measured in HEK293T cells at 24 h with the percentages of GFP^+^ cells at 48 h (**Figure 8d**), time point corresponding to the maximal fluorescence intensities for GFP^+^ and Cy5^+^, respectively. This dot plot further evidenced that the very low fluorescence in Cy5^+^ cells transfected with *b-*PEI_0.8_/Cy5-peGFP-C3 polyplexes did not allow any GFP expression. On the other hand, HEK293T cells transfected with *l*-PEI_20_/Cy5-peGFP-C3 polyplexes showed higher Cy5^+^ fluorescence means than those measured in cells incubated with *b*-PEI_60_/Cy5-peGFP-C3 polyplexes, while percentages of GFP^+^ cells were not significantly different (**Figure 8d**). In contrast, transfections with *b*-PEI_25_/(Cy5)-peGFP-C3 polyplexes led to higher Cy5^+^ fluorescence values correlating with higher percentages in GFP^+^ cells. The cells transfected with the lipofectamine 3000/Cy5-peGFP-C3 lipoplexes exhibited intermediate Cy5^+^ fluorescence means values between those incubated with *l*-PEI_20_/ and *b*-PEI_25_/(Cy5)-peGFP-C3 polyplexes, also correlated to an intermediate percentage of GFP^+^ cells.

**Figure 8.-.**
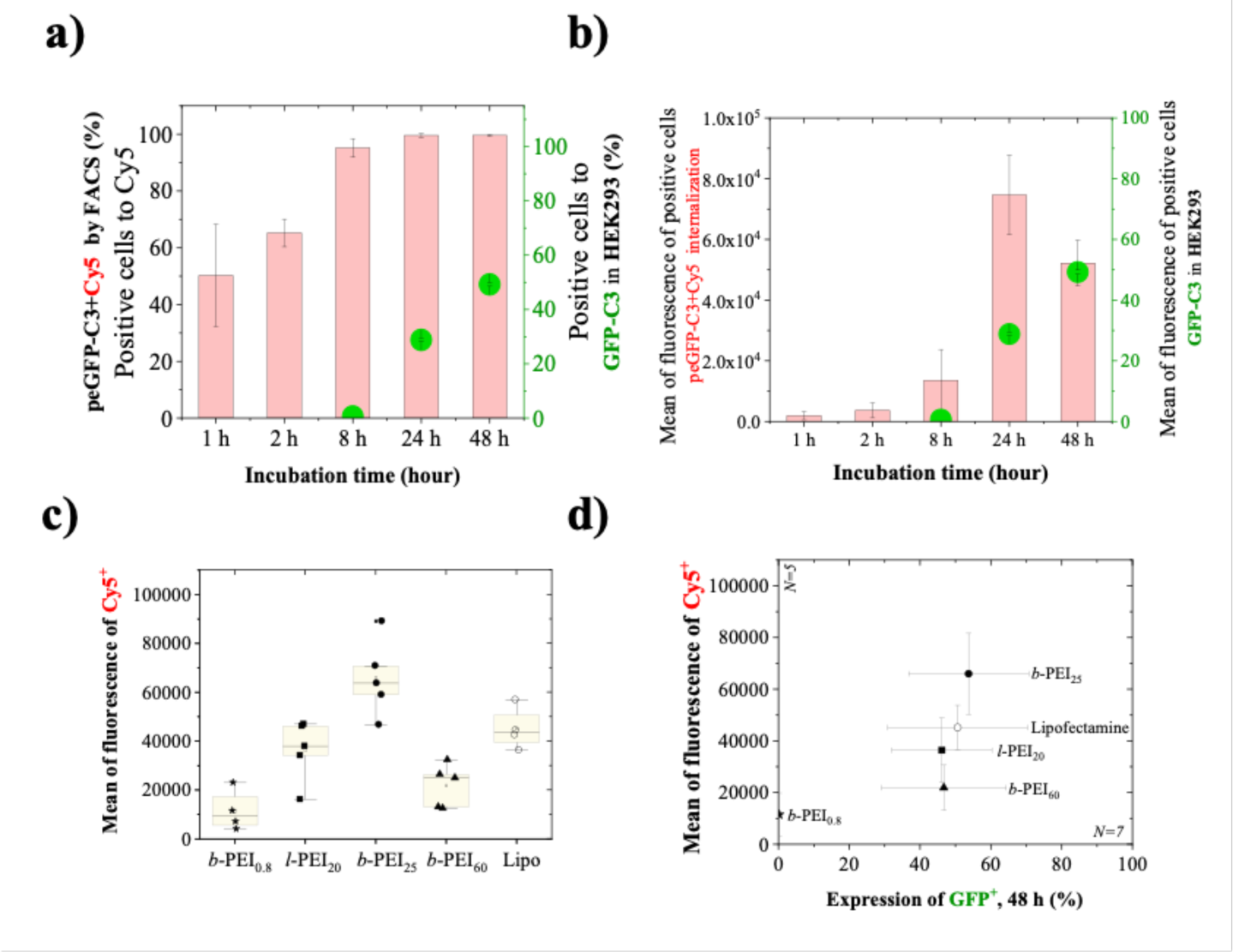
Quantification of flow cytometry data presenting (**a**) the percentages of positive HEK293T cells, and (**b**) their means of fluorescence transfected with *b*-PEI_25_/cyanine 5-peGFP-C3 polyplexes R=5 (left axis: Cy5^+^ cells) and cells transfected with *b*-PEI_25_/peGFP-C3 polyplexes R=5 (right axis: GFP^+^ cells) at 1, 2, 8, 24 and 48 hours after transfection. (**c**) Box plot presenting the means of fluorescence in cells transfected with with *b*-PEI_0.8_, *l*-PEI_20_, *b*-PEI_25_ and *b*-PEI_60_)/cyanine 5-peGFP-C3 (R=5) polyplexes and lipofectamine/cyanine 5-peGFP-C3 lipoplexes from 5 independent experiments. (**d**) Correlation graph presenting the means of fluorescence in cells transfected with with *b*-PEI_0.8_, *l*-PEI_20_, *b*-PEI_25_ and *b*-PEI_60_)/cyanine 5-peGFP-C3 (R=5) polyplexes and lipofectamine/cyanine 5-peGFP-C3 lipoplexes versus the percentage of GFP^+^ cells transfected with PEI/ and lipofectamine/peGFP-C3 encoding GFP.

In a final series of experiments, we addressed the question of the cellular localization of the different polyplexes and lipoplexes using confocal microscopy. The *b*-PEI_0.8_, *l-*PEI_20_, *b*-PEI_25_ and *b*-PEI_60_/Cy5-peGFP-C3 polyplexes and Lipofectamine 3000/Cy5-peGFP-C3 lipoplexes were detected by confocal microscopy to visualize their intracellular localization (**Figure 9, Figure S.I. 7**).

**Figure 9.-.**
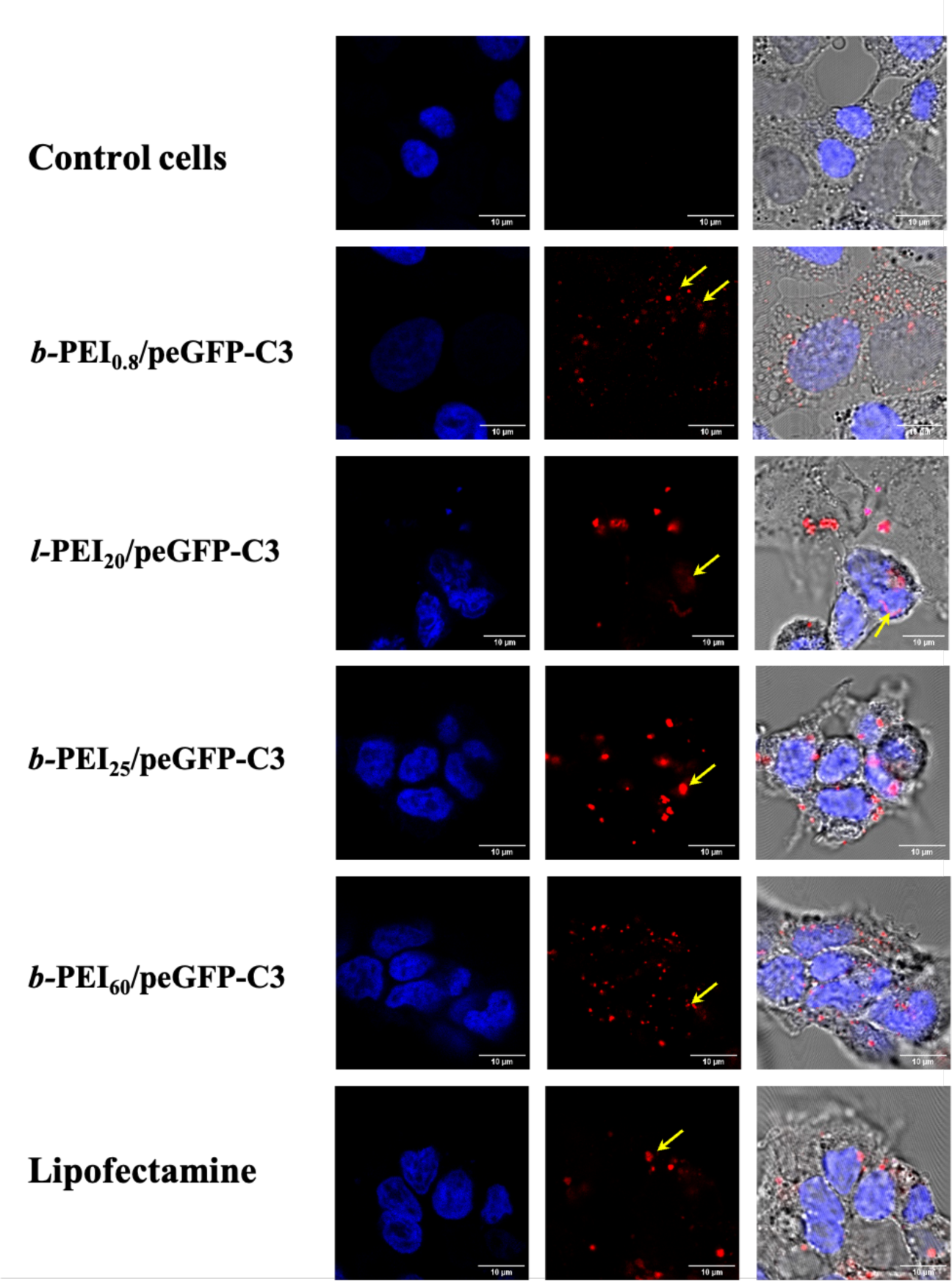
Detection of *b*-PEI_0.8_, *l*-PEI_20_, *b*-PEI_25_ and *b*-PEI_60_/ cyanine 5-peGFP-C3 polyplexes and lipofectamine/cyanine 5-peGFP-C3 lipoplexes (red fluorescence) in HEK293T cells with nuclear DNA stained with Hoechst 33342 (blue fluorescence). Yellow arrows indicate intracellular polyplexes and lipoplexes within the cytoplasm and near the nuclei. Control cells were non-transfected cells.

At low magnification, the observation of Cy5^+^ cells further confirmed that higher amounts of *l*-PEI_20_/(Cy5)-peGFP-C3 and *b*-PEI_25_/(Cy5)-peGFP-C3 polyplexes and Lipofectamine 3000/(Cy5)-peGFP-C3 lipoplexes were internalized compared to the lower fluorescence signals found in cells transfected with *b*-PEI_0.8_/(Cy5)-peGFP-C3 and *b*-PEI_60_/(Cy5)-peGFP-C3 polyplexes, as previously determined by flow cytometry. However, in all cells transfected with PEI-polyplexes or Lipofectamine-lipoplexes, intense fluorescent red speckles were observed inside the cells with an apparent random distribution.

At high magnification (**Figure 10**), in cells transfected with the *b*-PEI_25_/(Cy5)-peGFP-C3 polyplexes, the numerous fluorescent speckles present in the cytoplasm were either clearly distant from the nuclei or in close contact with the DNA stained with the blue dye Hoechst 33342 indicating that some of the plasmids reached the vicinity of the nucleus. In some cells, multiple speckles were found tightly associated with the stained nuclei strongly suggesting that plasmids had reached the nuclear compartment, in agreement with the fact that these *b*-PEI_25_/(Cy5)-peGFP-C3 polyplexes allow high GFP expression levels.

**Figure 10.-.**
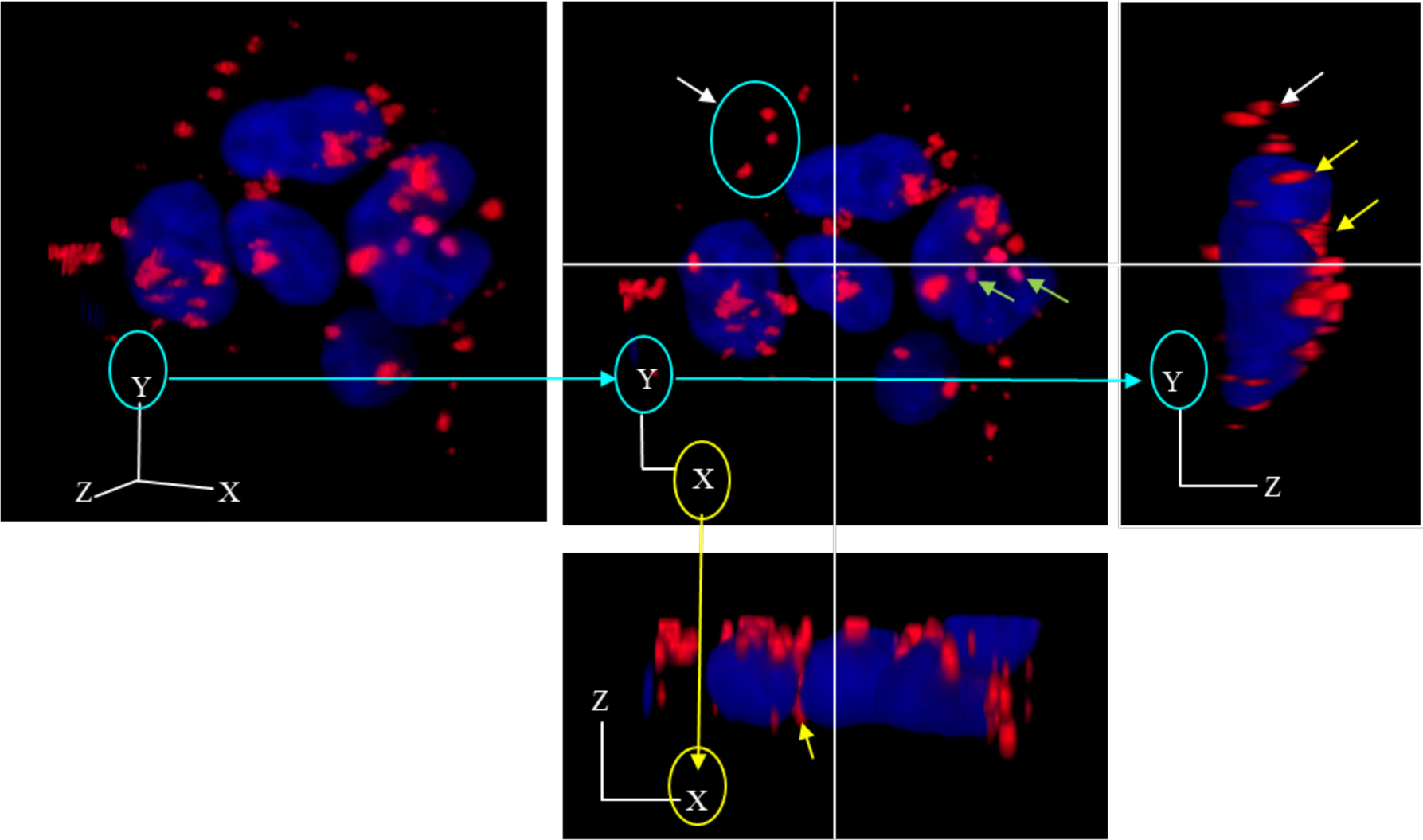
Detection of *b*-PEI_25_/cyanine 5-peGFP-C3 polyplexes (red fluorescence) in HEK293T cells with nuclear DNA stained with Hoechst 33342 (blue fluorescence). White arrows indicate intracellular polyplexes distant from the nuclei, yellow arrows point out polyplexes located near the nuclei and green arrows indicate polyplexes stained in pink probably resulting to merged speckles of polyplexes embedded in the blue-stained chromatin.

## 4. Conclusions

The ζ-potential values at the full complexation state of peGFP-C3 chains, corresponding to the highest ζ-potential for each PEI sample, were found to be dependent on PEI Mw. Full charge neutralization of peGFP-C3 with PEI was identified through the isoelectric point of PEI/peGFP-C3 polyplexes, occurring at a ζ-potential of 0 mV. At 2 ≤ R ≤ 15, ζ-potential of *b*-PEI_0.8_/peGFP-C3 polyplexes was ca + 20 mV, while ζ-potentials were significantly higher for *l*-PEI_20_, *b*-PEI_25_ and *b*-PEI_60_/peGFP-C3 polyplexes in the range of + 20 to + 40 mV, and increased with higher R. From AFM measurements, it was possible to observe particles with sizes around 50 nm and to identify the existence of a hierarchical complex structure, where small primary polyplexes formed during the initial phases of complexation aggregate to create bigger secondary polyplex particles. SAXS analysis allowed obtaining information about the sphere radius of the primary polyplexes, corresponding to be 19.7 ± 8.9 nm for *b*-PEI_0.8_/peGFP-C3, 7.9 ± 5.5 nm for *b*-PEI_25_/peGFP-C3 and 9.1 ± 5.4 nm for *b*-PEI_60_/peGFP-C3.

Our findings also revealed that PEI with too low Mw (*b*-PEI_0.8_) presents lower stability in presence of biological media, poor cell internalization and did not show any peGFP-C3 expression. We indeed studied the evolution of polyplexes size in PBS) and culture medium in absence and presence of serum. These later biological matrices constitute the real conditions for transfection experiments, however, stability of polyplexes and lipoplexes are rarely studied in these matrices. We found that *b*-PEI_0.8_/peGFP-C3 polyplexes not only showed poor stability in culture media especially at low charge ratios such as 3. Additionally, *l*-PEI_20_, *b*-PEI_25_ and *b*-PEI_60_/peGFP-C3 polyplexes displayed greater stability in culture media, particularly at R > 3. This finding supports the transfection data since we observed that both linear and branched PEI, with Mw of 20 and 25 kg/mol, respectively, at R ≥ 3, presented the highest levels of internalized plasmid and the best transfection efficiencies. However, polyplexes prepared with linear PEI displayed lower stability in presence of biological media compared to polyplexes prepared using branched PEI, with no plasmid DNA release even at low R. The relation between the % of GFP^+^ cells and the amounts of internalized plasmid revealed that *b*-PEI_25_ promoted higher internalization and peGFP-C3 expression on HEK293T cells. Herein, we also report that *b*-PEI_0.8,_ *l*-PEI_20_, *b*-PEI_25_ and *b*-PEI_60_/peGFP-C3 polyplexes and lipofectamine/peGFP-C3 lipoplexes are rapidly internalized in cells, which all contained plasmids within 8 hours after transfection. However, the overall amounts of *l*-PEI_20_ and *b*-PEI_25_/peGFP-C3 polyplexes progressively accumulated in HEK293T cells during 24 hours demonstrating the very dynamic process of internalization. In contrast, despite the fact that intracellular plasmids were visualized, GFP was not expressed in cells transfected with *b*-PEI_0.8_/peGFP-C3 polyplexes, suggesting that a threshold of plasmid amount must be achieved to allow protein expression. Our results highlight the impact of physicochemical parameters, some directly derived from PEI’s chemical structure, as well as on the excess of PEI for R>1, on polyplexes particle size, stability, cell internalization, DNA release, incubation time, cell viability and transfection efficiency. This study offers significant insights for selecting appropriate transfection conditions for efficient and safe for gene delivery. The obtained information represents the basis of further research aiming to study the dissociation mechanisms of PEI/peGFPC-3 polyplexes in a biological environment.

## Acknowledgements

The authors of this work acknowledge Bertrand Lefeuvre from the Institute of Chemical Sciences (ISCR) of Rennes for th<u>e</u> Malvern Zetasizer NanoZS facilities, Stephanie Dutertre (MRic) and Alexis Aimé (flow cytometry) core facilities of the Biology and Health Federative research structure Biosit (UAR 3480 CNRS – US18 Inserm), Rennes, France. Paulina Alejandra Montaño González acknowledges the Mexican National Council of Humanities, Sciences and Technologies, CONAHCYT (CVU 770728), the Organic Polymer Chemistry Laboratory (LCPO, Bordeaux), the Centre of Research Paul Pascal (CRPP, Bordeaux) and the Institute of Chemical Sciences (ISCR) of Rennes for technical facilities to carry on the research. Lourdes Mónica Bravo-Anaya acknowledges the ANR grant (ANR-21-CE06-0027-01), as well as the AES 2021 and the AIS Allocation 2023 founded by Rennes Métropole. This work was also funded by the Institut National de la Santé et de la Recherche Médicale (Inserm).

## Notes

### Competing Interest Statement

The authors have declared no competing interest.

