## Supplementary information for "Insight Into the Molecular Parameters of PEI Promoting an Efficient Gene Delivery into Cells"

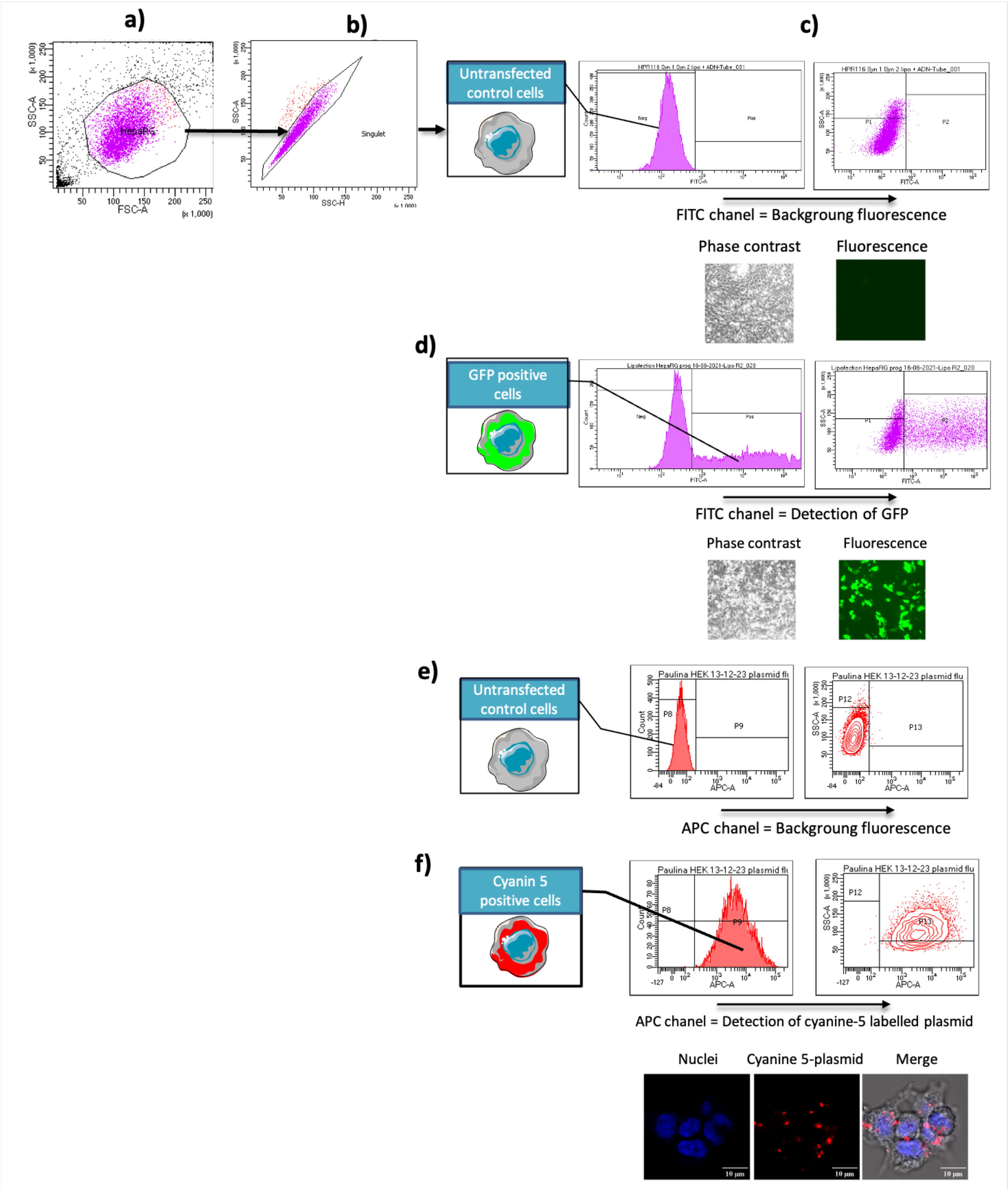


**Figure S.I. 1.-** The events (cells) analyzed by FACS were gated using dot plots with the Side SCatter (SSC, granularity on y axis) and Forward SCatter (FSC, size on x axis) parameters (**a**). Then, events from this first cell population were further gated using the SSC-Height versus SSC-Area dot plot to isolate the single cells (**b**). From the single cell population, fluorescence intensities were analyzed (histogram: fluorescence on x axis versus cell count on y axis; dot plot: fluorescence on x axis versus SSC-Area on y axis) in cells that were not transfected to define the "background" fluorescence also called auto-fluorescence of negative cells (Neg) on FITC (**c**) and APC (**e**) channels. In non-transfected control cells, no $GFP^+$ or cyanine $5^+$ cells were detected and the autofluorescence value (MFI) was arbitrarily set as 30 arbitrary units. Then, the cells transfected using plasmid encoding the GFP (**d**) or plasmid labeled with cyanine 5 (**f**) were analyzed, which defined the GFP and cyanine 5 positive cells (Pos). The values of MFI presented in this work represent the overall fluorescence of all the single $GFP^+$ cells (Pos). After acquiring fluorescence signals of ~$10^4$ cells, percentage of $GFP^+$ cells, fluorescence intensities of single cells were obtained from the CellQuest software. GFP positive ($GFP^+$) and cyanine 5 positive cells can be visualized by fluorescence microscopy.

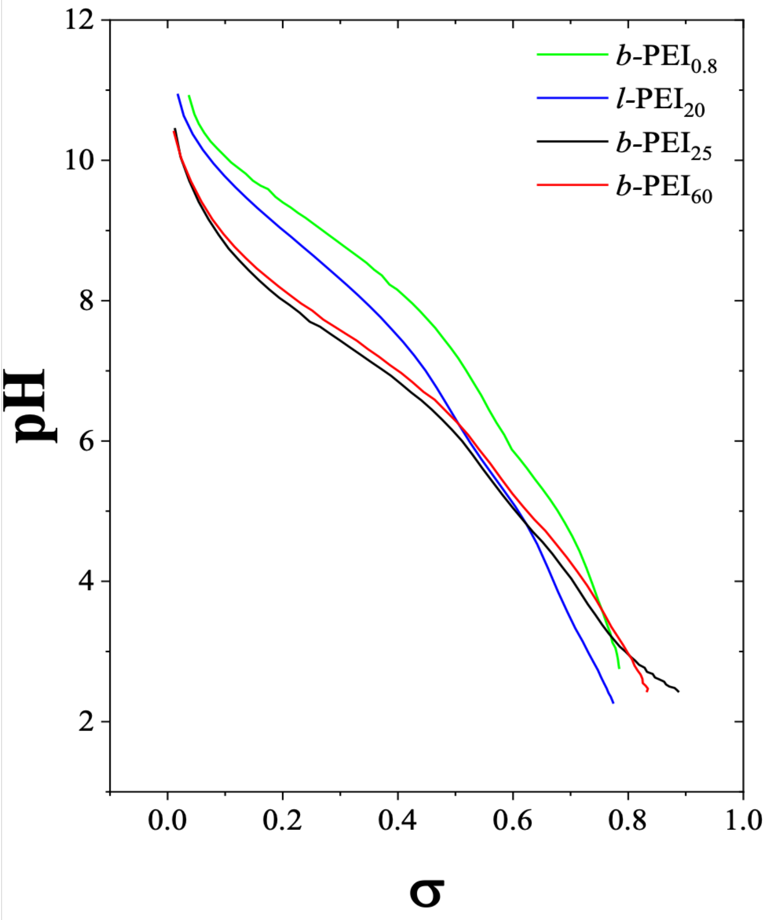


**Figure S.I. 2.-** Protonation rates as a function of pH calculated from the titrations of *b*-$PEI_{0.8}$, *b*-$PEI_{25}$ and *b*-$PEI_{60}$ (1 mg/mL, or 23 mM) with a 0.1 M HCl solution, and *l*-$PEI_{20}$ (1 mg/mL, or 12 mM) with a 0.1 M NaOH solution [1]. All of the titrations were carried on at 25 °C. The concentration of protonated amine groups, $[NH^+]$, including primary, secondary and tertiary amines was calculated by applying electroneutrality, $[NH^+] = [OH^-] + [Cl^-] - [H^+]$. The protonation rate (σ) was defined by $\sigma = [NH^+]/[N]^0$, where $[N]^0$ corresponds to the total amine concentration, determined from the molar concentration of repetitive units in *b*-$PEI_{25}$ and *l*-$PEI_{20}$.


*[1] C. J. B. van Treslong, A. J. Staverman, Poly (ethylenimine) II. Potentiometric titration behaviour in comparison with other weak polyelectrolytes, Recl. Trav. Chim. Pays-Bas 93 (6) (1974) 171–178. https://doi.org/10.1002/recl.19740930612.*

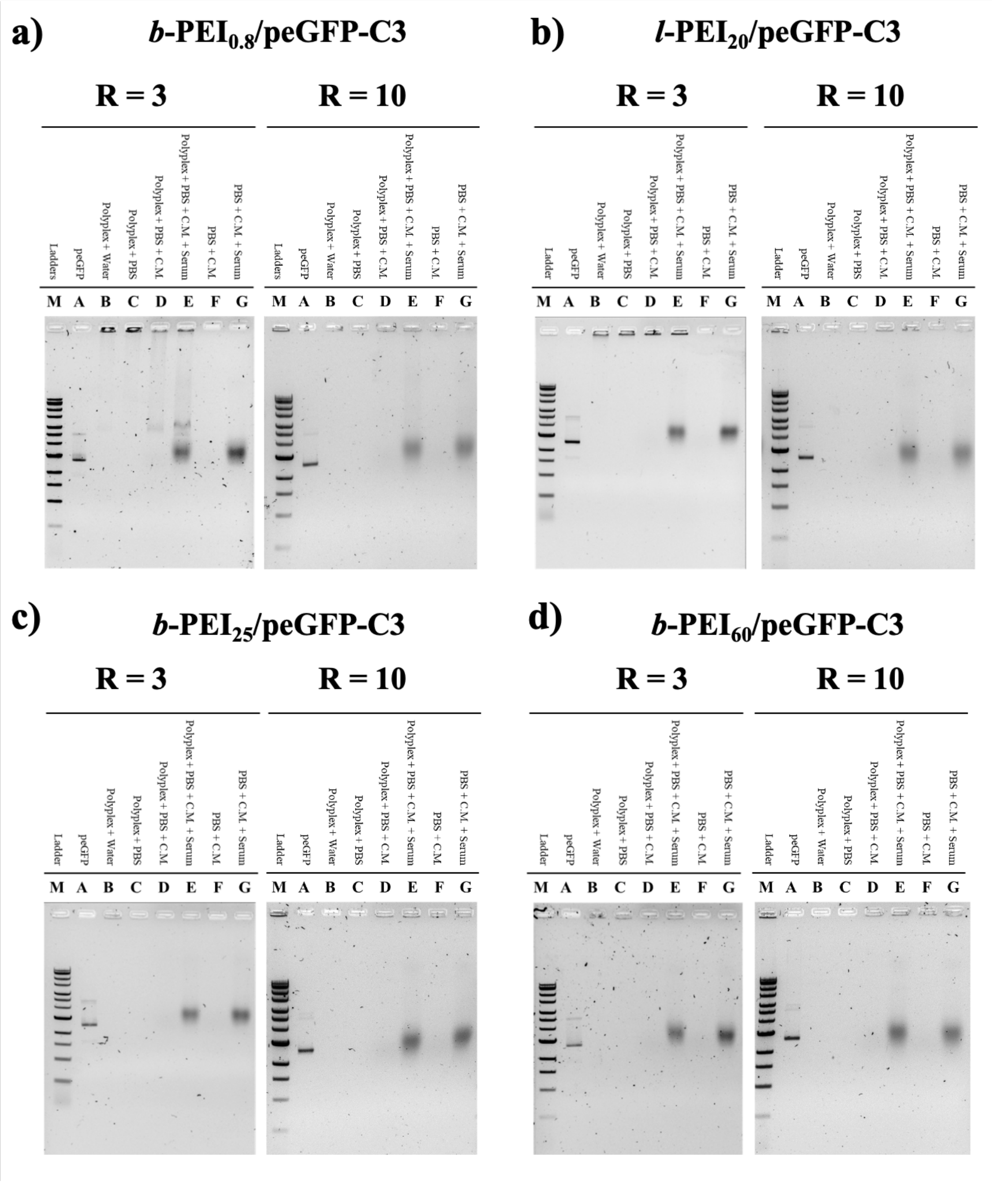


**Figure S.I. 3.-** Electrophoresis gel assays showing the colloidal stability, in terms of DNA release, of PEI/peGFP-C3 polyplexes prepared at two different charge ratios ($[N^+]/[P^-]$= 3 and 10), using: a) *b*-$PEI_{0.8}$, b) *l*-$PEI_{20}$, c) *b*-$PEI_{25}$ and d) *b*-$PEI_{60}$, in presence of PBS, culture media (CM) and culture media supplemented with serum. M represents the molecular weights of the marker (SmartLadder MW-1700-10 Eurogentec). $C_{peGFP\text{-}C3}$ = 0.034 mg/mL in water, $C_{b\text{-}PEI0.8}$ = 0.8925 mg/mL, $C_{l\text{-}PEI20}$ = 0.8721 mg/mL, $C_{b\text{-}PEI25}$ = 0.9295 mg/mL and $C_{b\text{-}PEI60}$ = 0.9254 mg/mL in water (pH 7.4). Samples containing 15.84 ng pDNA were loaded onto a 0.8 % agarose gel (final concentration: 0.022 mg/mL of peGFP-C3).

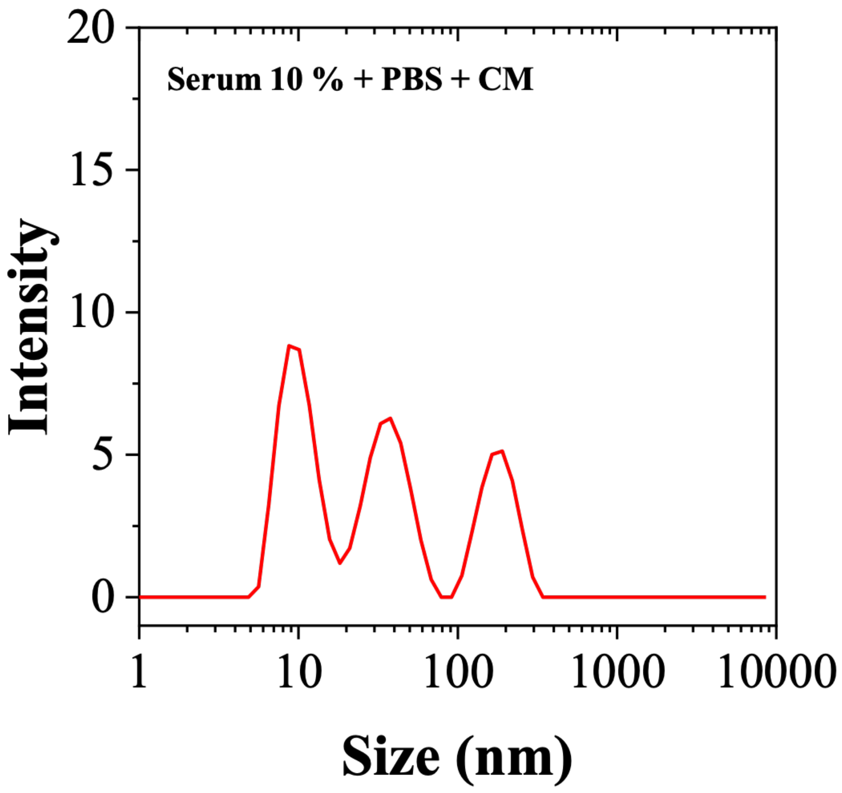


**Figure S.I. 4.-** Intensity-averaged size distribution obtained by DLS for PBS and DMEM supplemented with 10% fetal bovine serum (FBS). Measurements were performed at 25 °C.

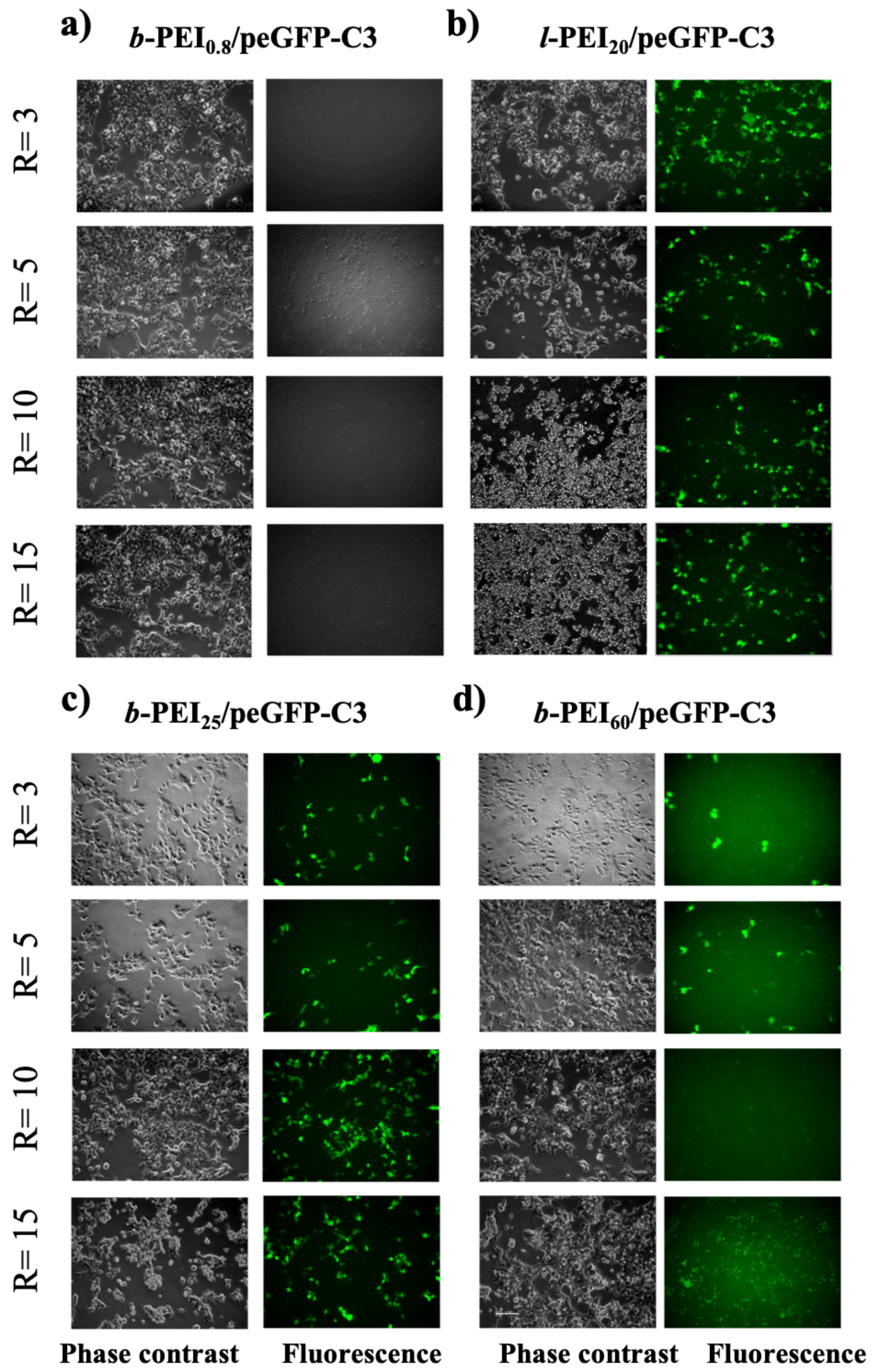


**Figure S.I. 5.-** The HEK293T cells transfected with *b*-$PEI_{0.8}$, *l*-$PEI_{20}$, *b*-$PEI_{25}$ and *b*-$PEI_{60}$/peGFP-C3 polyplexes, prepared at four charge ratios (R = $[N^+]/[P^-]$ = 3, 5, 10 and 15), were observed in phase contrast and fluorescence microscopy. GFP positive cells can be visualized by fluorescence microscopy (U.V. excitation).

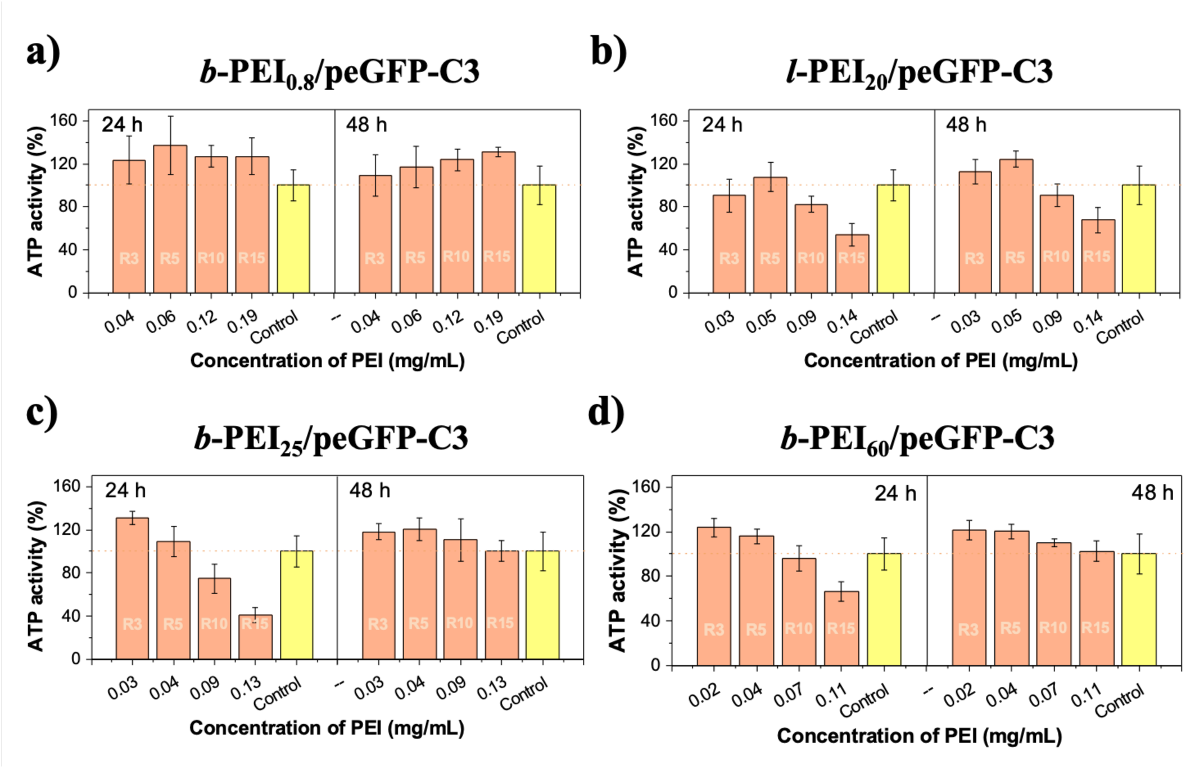


**Figure S.I. 6.-** The cell viability was assayed in HEK293T cells transfected with *b*-$PEI_{0.8}$, *l*-$PEI_{20}$, *b*-$PEI_{25}$ and *b*-$PEI_{60}$/peGFP-C3 polyplexes at four charge ratios (R = $[N^+]/[P^-]$ = 3, 5, 10 and 15), corresponding to various concentrations in the corresponding PEI (expressed in mg/mL). Cell viability was measured with the relative ATP content reflecting the number of cells. Results were expressed in percentage of the cell viability in non-transfected control cells arbitrary set as 100%. While transfection with *b*-$PEI_{0.8}$/peGFP-C3 polyplexes had no significant effect on cell numbers, transfection with *l*-$PEI_{20}$, *b*-$PEI_{25}$ and *b*-$PEI_{60}$/peGFP-C3 polyplexes reduced the cell number in a concentration dependent manner (at high R = $[N^+]/[P^-]$ = 10 and 15) at 24 hours after incubation. This decreased number of cells compared to control culture condition may be due in part to cell death but also to a cytostatic effect since after discarding the transfection media at 24 hours, the decrease in cell number was no longer observed at 48 hours.

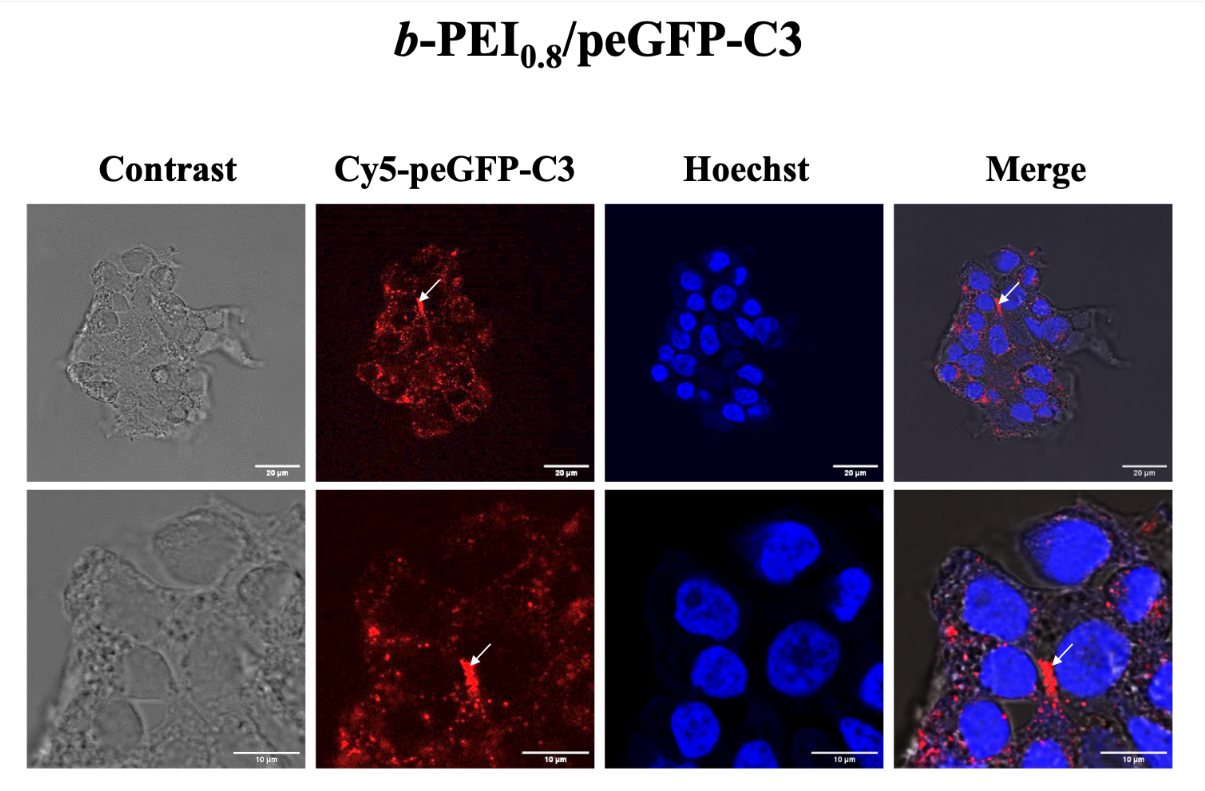


**Figure S.I. 7.-** Detection of *b*-PEI$_{0.8}$/cyanine 5-peGFP-C3 polyplexes (red fluorescence) in HEK293T cells with nuclear DNA stained with Hoechst 33342 (blue fluorescence). The lower panels are high magnifications of the upper photographs.
